# A computational and structural analysis of germline and somatic variants affecting the DDR mechanism, and their impact on human diseases and prostate cancer progression

**DOI:** 10.1101/2021.01.21.427605

**Authors:** Lorena Magraner-Pardo, Roman A. Laskowski, Tirso Pons, Janet M. Thornton

**Affiliations:** Prostate Cancer Clinical Unit, Spanish National Cancer Research Center (CNIO), Madrid, Spain; European Molecular Biology Laboratory, European Bioinformatics Institute (EMBL-EBI), Cambridge, UK; Department of Immunology and Oncology, National Center for Biotechnology, Spanish National Research Council (CNB-CSIC), Madrid, Spain

## Abstract

DNA-Damage Response (DDR) proteins are crucial for maintaining the integrity of the genome by identifying and repairing errors in DNA. Variants affecting their function can have dire consequences as damaged DNA can result in cells turning cancerous. Here we compare germline and somatic variants in DDR genes, specifically looking at their locations in the corresponding three-dimensional (3D) structures, Pfam domains, and protein-protein interaction interfaces. We show that somatic variants are more likely to be found in Pfam domains and protein interaction interfaces than are pathogenic germline variants or variants of unknown significance (VUS). We also show that there are hotspots in the structures of ATM and BRCA2 proteins where pathogenic germline, and recurrent somatic variants from primary and metastatic tumours, cluster together in 3D. Moreover, in the *ATM, BRCA1* and *BRCA2* genes from prostate cancer patients, the distributions of germline benign, pathogenic, VUS, and recurrent somatic variants differ across Pfam domains. Together, these results provide a better characterisation of the most recurrent affected regions in DDRs and could help in the understanding of individual susceptibility to tumour development.

## Introduction

DNA is subject to continuous damage-causing alterations in both somatic and germline tissues. Cells rely on their DNA-Damage Response (DDR) systems to repair the damage before it leads to serious harm, such as the cell turning cancerous. Different pathways are involved, depending on whether the DNA sequence damage is a mutated base, a single-strand break (SSB), or a double-strand break (DSB). These pathways include: Base Excision Repair (BER), Nucleotide Excision Repair (NER), Mismatch Repair (MMR), Homologous Recombination (HR) and Non-homologous End Joining (NHEJ). Each pathway has its own set of DNA-repair proteins, encoded by the DDR genes (Andrés-León et al., 2016; Knijnenburg et al., 2018). A proper coordination between the repair machinery, damage tolerance, and checkpoint pathways is required for preserving DNA integrity and proper cell function (Lans et al., 2019).

A list of 276 DDR genes was recently identified by The Cancer Genome Atlas (TCGA) DNA Damage Repair Analysis Working Group which systematically analysed potential causes of loss of DDR function across 33 different cancer types and their consequences in human cancer (Knijnenburg et al., 2018). This set of genes provides the basis for the mechanistic and therapeutic analysis of the role of DDR in cancer. In addition, pan-cancer studies provide evidence for factors affecting predisposition to different cancer types, highlighting rare germline cancer susceptibility variants that affect tumour suppressor genes including *ATM*, *BRCA1*, *BRCA2*, *BRIP1*, and *PALB2* (Lu et al., 2015). These and other studies looked at somatic and germline variants and their clinical associations to help understand individual susceptibility to tumour development (Mamidi et al., 2019; Sivley et al., 2018). However, a large structural analysis of candidate DDR genes accumulating pathogenic germline and recurrent somatic variants is not yet available.

Concerning prostate cancer (PCa), it has been reported that genetic variations in some DDR genes not only increase the risk of the disease, but are also associated with poor prognosis and clinical outcomes at different PCa stages (Edwards et al., 2008). Previous studies found that 22.7% (34/150) of metastatic, castration-resistant PCa (mCRPCa) patients harbour biallelic somatic/germline variants in DDR genes, and specifically 8% had germline variants (Robinson et al., 2015). Other studies found that 11.8% (82/692) of patients with mCRPCa had at least one germline variant in a gene involved in DNA repair (Pritchard et al., 2016). These and other mCRPCa-based studies (Annala et al., 2018; Antonarakis et al., 2018; Castro et al., 2019; Mijuskovic et al., 2018; Na et al., 2017; Pritchard et al., 2016; Robinson et al., 2015) highlighted *BRCA2* as the most frequently involved, and also found that a high proportion of those patients presented a second “hit” somatic aberration in either of two forms: loss-of-function mutation or a gene-copy-loss (Jonsson et al., 2019; Pritchard et al., 2016; Sokol et al., 2020). The same authors suggested that only half of the patients could be detected by germline variants in DDR genes. Based on these studies, there is an evident need to test or examine routinely all patients with mCRPCa for the presence of germline/somatic variants in DDR genes.

In this article, we study the different distributions of germline benign, pathogenic, variants of unknown significance (VUS), and recurrent somatic variants across Pfam domains, three-dimensional (3D) structures, and protein interaction interfaces in DDR genes. We find that the percentage of somatic variants accumulated in functional domains and protein interactions interfaces, is higher than in germline pathogenic variants and VUS, providing a combined effect to damage the protein function. In addition, for the ATM and BRCA2 proteins, we identified 3D clusters and hotspot regions that accumulate pathogenic germline and recurrent somatic variants from primary and metastatic tumours, showing the synergistic effect of a few mutations located in specific regions and contributing to oncogenesis. With this analysis, we provide a better characterisation of the most recurrent affected functional regions in DDRs, which can help the understanding of an individual’s susceptibility to tumour development.

## Results

### DDR genes accumulate more germline variants than their non-DDR interactors in the NetDDR network

We first used the NetDDR interaction network to investigate the importance of the DDR protein-coding genes in terms of their protein interactions and accumulation of genetic variants. It has been suggested that mutations in regulatory regions of highly connected protein-coding genes in protein-protein interaction and regulatory networks have a higher functional impact than those targeting peripheral genes in the network (Khurana et al., 2013). The number of variants studied in the different classes were: for germline variants (1) 10,301 pathogenic and likely pathogenic, (2) 1,117 benign and likely benign, and (3) 28,248 VUS, whereas for somatic variants (1) 5,795 variants in primary tumours and (2) 2,030 variants in metastasis.

We found that recurrent somatic variants (i.e. occurring in more than one sample) annotated in COSMIC are found in more than 93% of both DDR and non-DDR interacting proteins in the NetDDR network (Figure 1, panel A). However, germline variants annotated from ClinVar have been identified in DDR (40%) more than in non-DDR (21%) genes (Figure 1, panel B). This finding suggests there is some bias towards the most studied genes, human variations, and phenotypes with supporting evidence annotated in the ClinVar database whose primary source of information are published studies. In contrast, the COSMIC database documents somatic mutations in human cancers not only from published studies, but also from whole genome and exome sequencing experiments that identify variants more homogeneously across all human genes.

**Figure 1:**
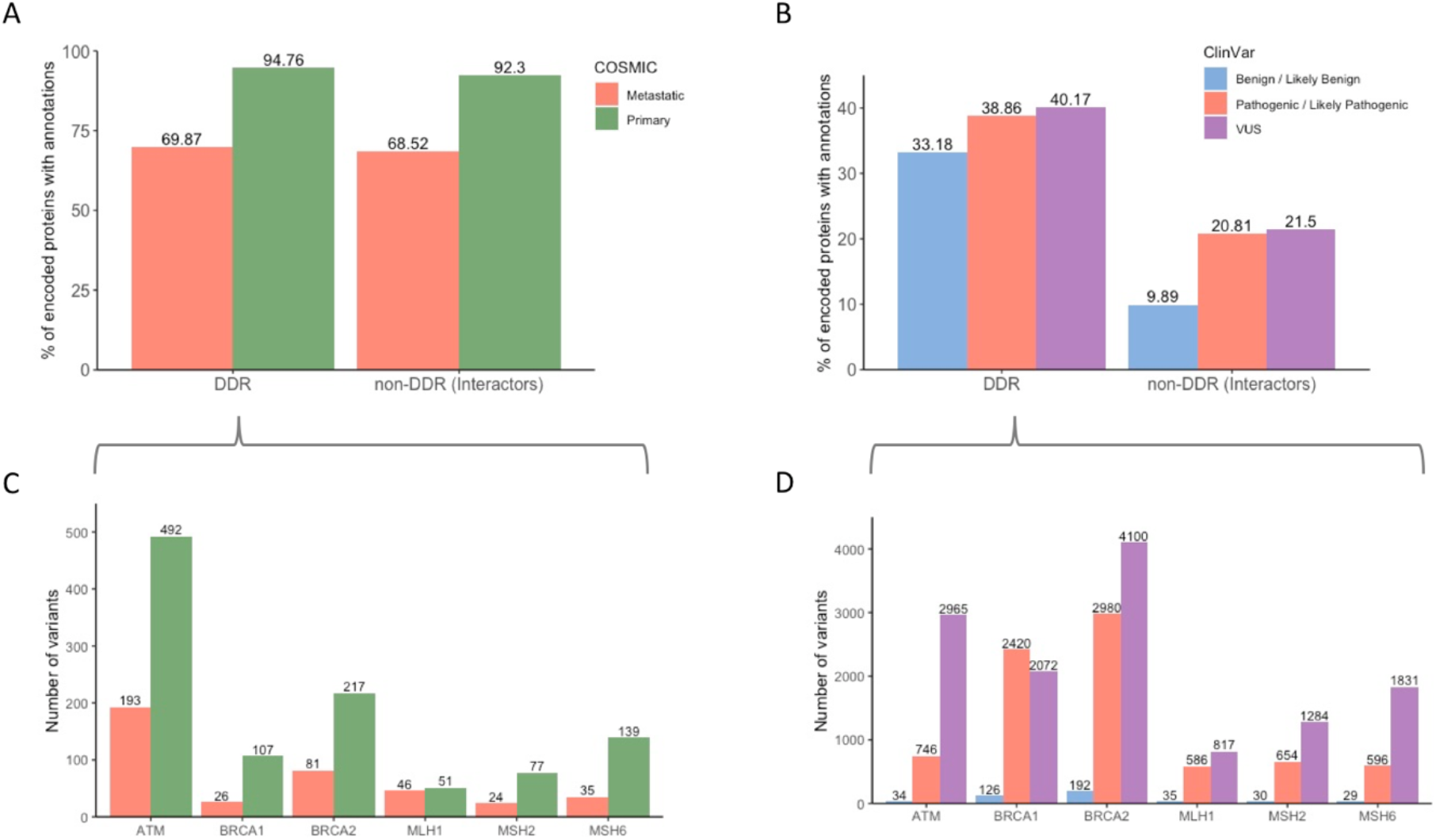
Accumulation of germline and recurrent somatic variants in DDR and non-DDR interactors in the NetDDR network. Percentages of protein-coding genes with variant annotations in COSMIC and ClinVar are shown in panels A and B, respectively. Patterns of recurrent somatic (≥2 samples) and germline variants in the highly mutated DDR genes are illustrated in panels C and D, respectively.

In addition, the largest set of ClinVar annotations are VUS variants, followed by pathogenic and benign, thus showing another example of bias in the dataset. Other authors have observed some of the biases associated with ClinVar, in particular, the inflated pathogenic variants profiles that make it difficult to study variant penetrance and disease prevalence in endocrine tumour syndromes (Toledo & Group, 2018). In the COSMIC database we found fewer variants identified in metastatic tumours than in a primary state, probably due to the low number of individuals studied with advanced cancer.

Also shown in Figure 1 are the numbers of somatic and germline variants in the cancer-predisposition genes *ATM*, *BRCA1*, *BRCA2*, *MLH1*, *MSH2*, and *MSH6* (Figure 1, panels C and D). These DDR genes are among the most affected in the COSMIC and ClinVar datasets (Supplementary Figure 5). The majority of recurrent somatic variants, observed in more than two samples, were from primary tumours. In addition, most germline variants in these cancer-predisposition and DDR genes correspond to pathogenic and VUS, although their ratios varied depending on the gene analysed. Besides, these cancer-predisposition genes are highly connected and central in the NetDDR interaction network. Indeed, the average node degree (AvrNodDeg) for these six genes is 166, while in the complete DDR-hits AvrNodDeg=87 and in non-DDR AvrNodDeg=45. Also, closeness centrality (ClossCentl), which indicates how close a node is to all other nodes in the network, is higher on average for the DDR-hits (ClossCentl=0.46) than for non-DDR (ClossCentl=0.42) (for more details see Supplementary Table 1). In the sections below we also investigated the effects of protein length, domain composition, and protein 3D structure on the accumulation of germline and somatic variants in DDR protein-coding genes.

### DDR genes shows a different pattern of accumulation of germline and somatic variants as a function of protein length

We analysed the distribution of germline and somatic variants normalized by protein length (Figure 2). The proteins in our dataset range in length from 44 a.a. (in TMSB4X) to 7,968 a.a. (OBSCN). The boxplot in Figure 2, panel A, shows that the mean number of germline variants per protein length in the DDR proteins is 0.287, which is significantly higher than the mean of 0.106 for the non-DDR (P-value = 5.0×10^−4^). Panel B shows the differences with the variants grouped by the different categories. A few DDR proteins appear above the 75th percentile, indicating accumulation of a high number of germline variants. These proteins are BRCA1, BRCA2, MSH2, MLH1, and MSH6 (Supplementary Figure 5).

**Figure 2:**
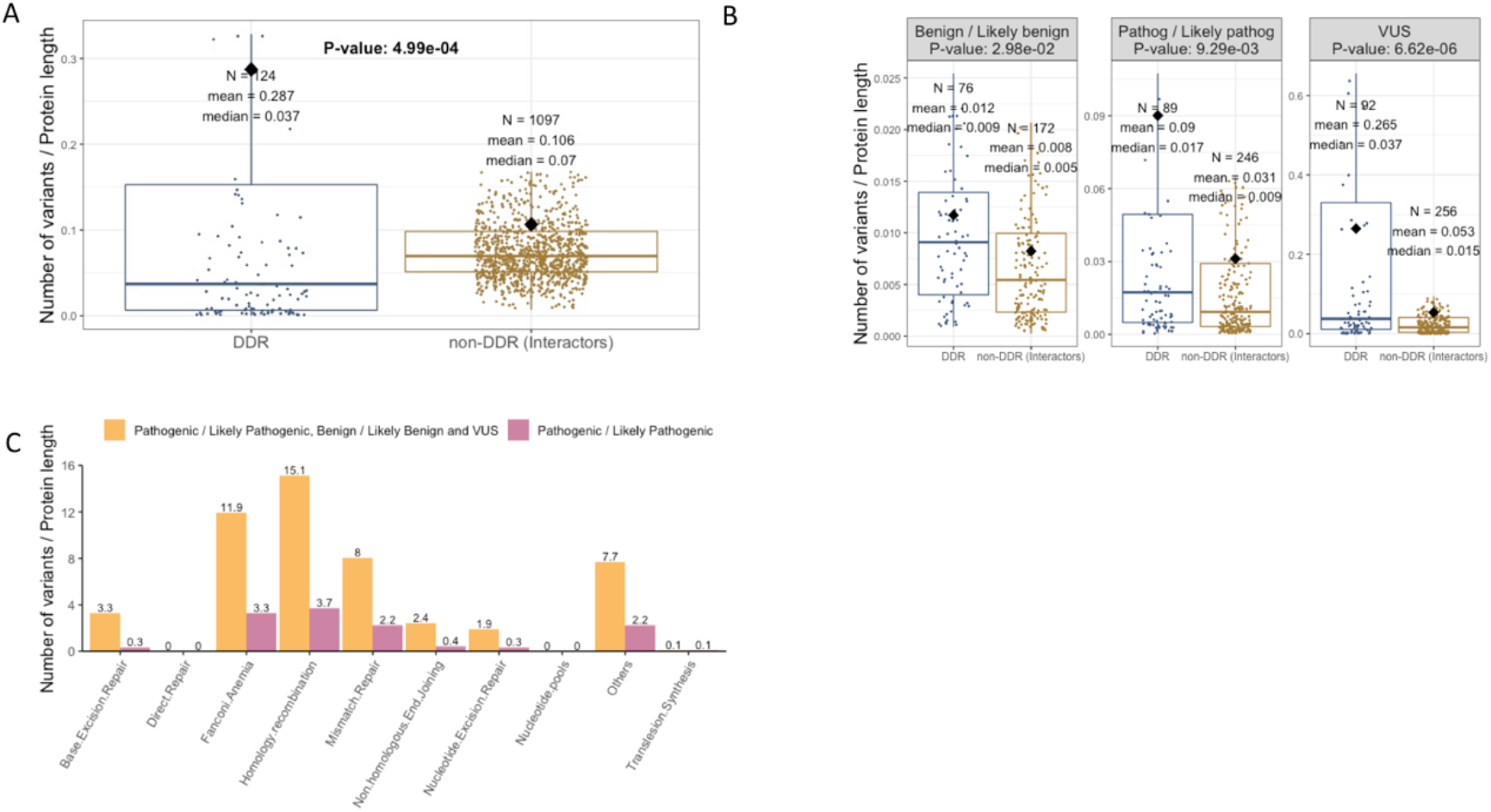
Distributions of germline variants in DDR and non-DDR interactors. Boxplots showing the contrasting patterns of germline variants as a function of protein length in the DDR and non-DDR proteins (panel A), and as categorized into benign, pathogenic, and VUS (panel B). All box plots depict the first and third quartiles as the lower and upper bounds of the box, with a thicker band inside the box showing the median value, and whiskers representing 1.5× the interquartile range. Panel C shows the accumulation of germline variants in biological pathways related to DDR.

Moreover, Figure 2, panel B, shows a greater variability in the DDR distributions and also shows that the medians (at 95% confidence interval) tend to be larger than for the non-DDR group, with P-values being 3.0×10^−2^ for germline benign, 9.3×10^−3^ for pathogenic, and 6.6×10^−6^ for VUS. Panel A shows a few DDRs above the 75th percentile, so we hypothesize that the accumulation of a large number of ClinVar pathogenic and VUS annotations in these DDRs (i.e. BRCA1, BRCA2, MLH1, MSH2) could bias research findings or limit the generalizability of the results.

In addition, we analysed the accumulation of germline and somatic variants in the different pathways where DDRs are involved. We found that the Homology Recombination, Fanconi Anaemia, and Mismatch Repair pathways are the most affected by germline mutations (Figure 2, panel C). The analysis of recurrent somatic variants annotated in COSMIC, also indicated the same affected pathways (Figure 3).

**Figure 3:**
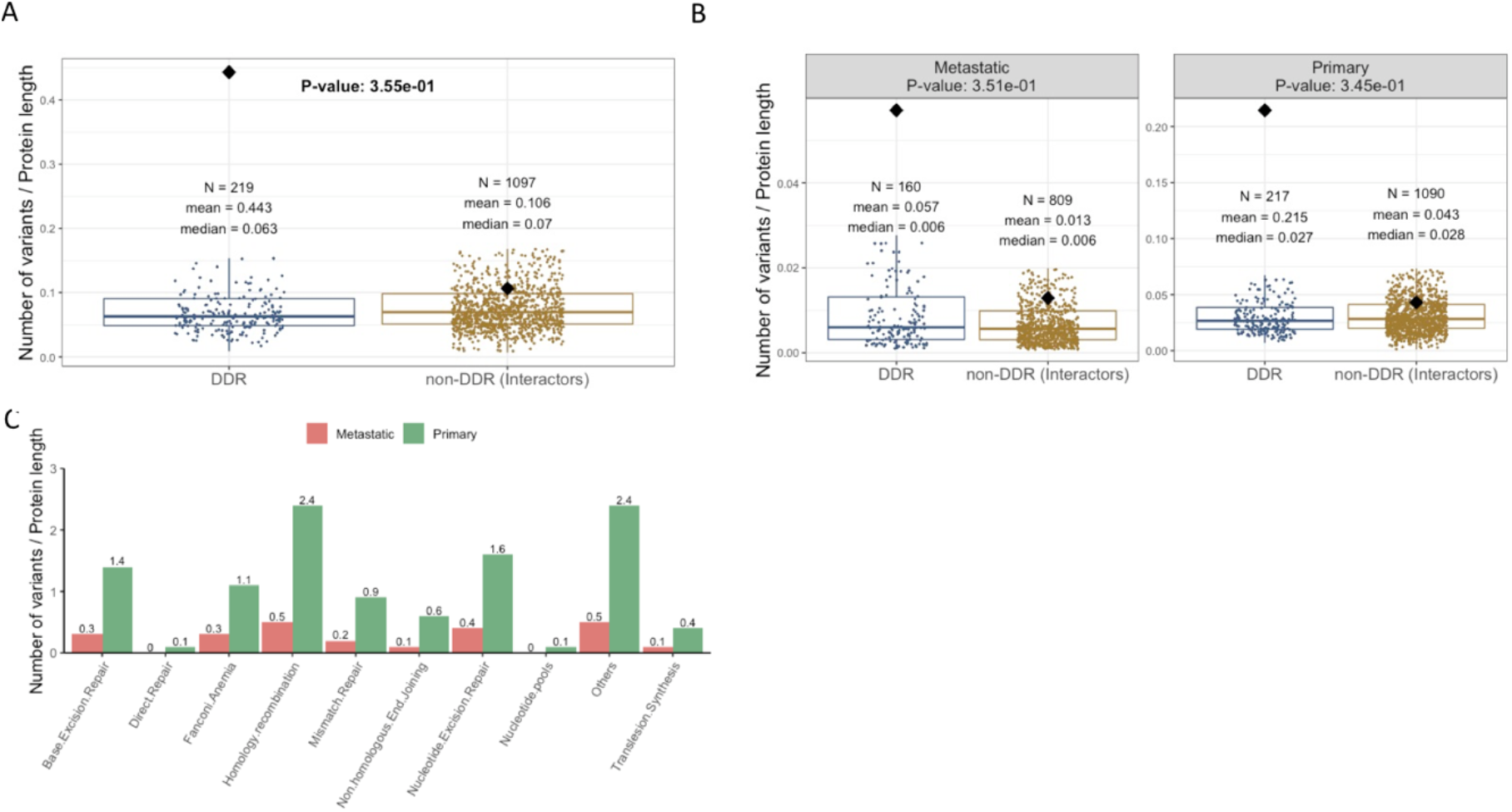
Distributions of somatic variants in DDR and non-DDR interactors. Boxplots showing the similar patterns in somatic variants and abundances in DDR and non-DDR (panel A) and for the somatic variants in metastasis and primary tumours (panel B). All box plots depict the first and third quartiles as the lower and upper bounds of the box, with a thicker band inside the box showing the median value and whiskers representing 1.5× the interquartile range. Panel C shows the accumulation of germline variants in different biological pathways associated with DDR genes.

The boxplots in Figure 3 (panels A and B) show similar medians and dispersion of the somatic variants per protein length in DDR and non-DDR groups (P-value = 0.36; metastatic: P-value = 0.35, primary: P-value = 0.35). However, this pattern of variability is different to the one previously observed in the germline variants annotated in ClinVar. In this case, *ATM*, *ATRX*, *CHEK2*, *ERCC2*, *MLH1*, *MSH6*, and *SMARCA4*, among other genes, are representative DDRs above the 75th percentile that indicate accumulation of somatic variants from the COSMIC database. Moreover, according to panel C in Figure 3, only the Nucleotide Excision Repair (NER) pathway appears to be more affected by somatic variants in primary tumours than by germline pathogenic variants. Therefore, our analyses of accumulation of somatic variants in the NER pathway coincide with the results by other authors in that this pathway shows an increased contribution of a somatic mutational pattern (COSMIC mutational signature 8) recurrently observed in various cancer types (Jager et al., 2019). So, the disruption of this pathway could potentially drive carcinogenesis and accelerate aging.

### DDR germline and somatic variants occur differently within Pfam domains and protein interaction interfaces

We also investigated the occurrence of DDR germline and somatic variants in Pfam domains and protein interaction interfaces. The curated list of germline and recurrent (≥2 samples) somatic variants were annotated using the Structure-PPi system (Vázquez et al., 2015) and the results are summarized in Figure 4 and in Supplementary Table 2.

**Figure 4:**
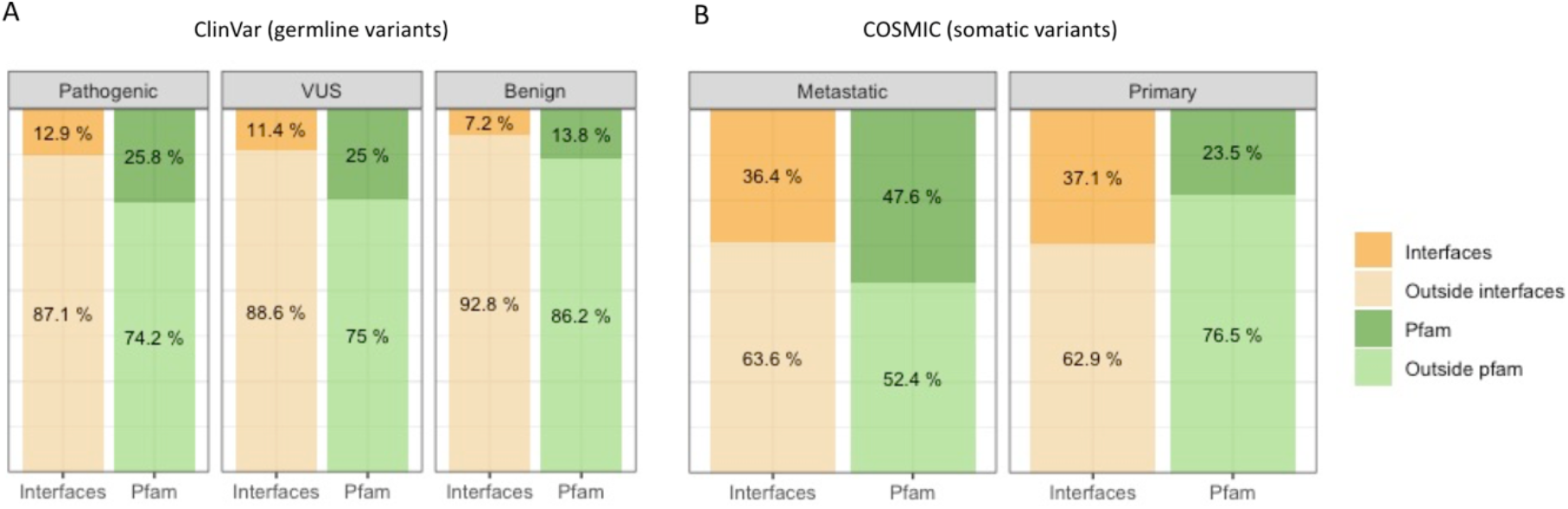
Accumulation of germline and somatic variants in Pfam domains and protein interaction interfaces. Panel A shows the percentage of pathogenic, VUS and benign germline variants extracted from the ClinVar database across interfaces and Pfam domains. Panel B shows the percentage of metastatic and primary somatic variants extracted from COSMIC across interfaces and Pfam domains.

These results indicate that germline variants in the pathogenic and VUS classes have a similar distribution (Figure 4, panel A). In particular, they possess a very similar percentage of variants mapped onto protein interaction interfaces (12.9% and 11.4%, respectively) higher than in the case of benign variants (7.2%). Interestingly, the percentage of somatic variants in metastatic tumours found in both Pfam domains and protein interactions interfaces, is higher than in germline pathogenic and VUS variants. The 47.6% metastatic variants affecting Pfam domains are double the value of 25% for germline variants. Otherwise, the percentage of metastatic variants (36.4%) affecting protein interaction interfaces is 3-fold higher than the value of 11% to 13% in germline variants.

The most affected Pfam domains by germline pathogenic variants were: MutS_III (Pfam code: PF05192, 371 variants), MutS domain V (Pfam code: PF00488, 323 variants), and BRCA2 (Pfam code: PF00634, 224 variants). MutS domains are found in human MSH2/MSH6 proteins implicated in non-polyposis colorectal carcinoma (HNPCC), while *BRCA2* is a known tumour suppressor gene. For the BRCA2 or FANCD1 proteins, their association with the Fanconi anaemia (FANC) protein complex is well known (D’Andrea & Grompe, 2003). Other Pfam domains affected with germline pathogenic variants are: BRCA2-helical (Pfam code: PF09169, 159 variants), BRCA2-OB1 (Pfam code: PF09103, 116 variants), PTEN-C2 (Pfam code: PF10409, 110 variants), BRCA1 C-terminus (Pfam code: PF00533, 108 variants), P53 (Pfam code: PF00870, 107 variants) and MutS domain IV (Pfam code: PF05190, 105 variants). It is very interesting that the DDR pathogenic germline variants in functional domains tend to occur in the known tumour suppressors *P53*, *BRCA2* and *PTEN*. But as we commented before, this observation would suggest some bias towards the most studied genes and phenotypes with supporting evidence annotated in the ClinVar database.

On the other hand, Pfam domains affected by somatic variants identified in metastatic tumours also included the known tumour suppressor genes *P53* and *PTEN*: P53 DNA-binding (Pfam code: PF00870, 408 variants), PTEN-C2 (Pfam code: PF10409, 44 variants) and P53 tetramer (Pfam code: PF07710, 27 variants). A description of the affected Pfam domains discussed in this article and the associated functions is presented in Table 4.

Overall, these observations agree with the hypothesis of two-hit events: first a germline variant produces a predisposition to develop a tumour, while a second somatic event increases this probability in a specific organ and triggers tumour development.

### Co-localization of pathogenic germline and somatic variants in protein sequence and 3D structure in DDR genes ATM, BRCA1, BRCA2, and MUTYH

In a previous work, we described *ATM*, *BRCA1, BRCA2*, and *MUTYH* as recurring genes with mutations in the PROREPAIR-B PCa cohort (Castro et al., 2019). Here, as use cases in the DDR study, we studied these genes and analysed more pathogenic germline variants identified in different cohorts of advanced PCa reported in the literature (**Table 1**), recurrent somatic and germline variants in PCa as well as hotspot positions in different tumour types collected from cBioPortal (https://www.cbioportal.org) (Supplementary File 1), and also, the TCGA-PanCancer study of pathogenic germline variants in 10,389 adult cancers (K. lin Huang et al., 2018).

**Table 1:** Pathogenic germline variants identified in mCRPCa cohorts and affecting the PCa relevant genes ATM, BRCA1, BRCA2 and MUTYH. (*) Numbers indicate the total variants identified in each study and in parenthesis the non-identical germline variants. (a) From 11 PCa studies in the non-redundant dataset at cBioPortal. (b) From 176 different studies in the non-redundant dataset at cBioPortal. Details about the PCa populations in cBioPortal are provided in Supplementary File 1.

| **Publication** | ***ATM*\*** | ***BRCA1*\*** | ***BRCA2*\*** | ***MUTYH*\*** | **Population (n)** | **%*BRCA2***<br>**mutated** |
| --- | --- | --- | --- | --- | --- | --- |
| Robinson et al. 2015 | 8 | 4 | 11 | - | 150 | 7.3 |
| Pritchard et al. 2016 | 11 | 6 (3) | 37 (24) | - | 692 | 5.35 |
| Na et al. 2017 | 6 | 2 | 11 | - | 313 | 3.51 |
| Mijuskovic et al. 2018 | 3 | 0 | 4 | - | 139 | 2.88 |
| Annala et al. 2018 | 1 | 1 | 16 (13) | - | 202 | 7.92 |
| Antonarakis et al. 2018 | 3 | 1 | 5 (3) | - | 172 | 2.91 |
| PROREPAIR-B<br>(Castro et al., 2019) | 8 (5) | 4 | 14 (13) | 13 (3) | 419 | 3.34 |
| cBioPortal (recurrent) | 10 | 2 | 16 | 1 | 1,556 <sup>(a)</sup> | 8.0 |
| cBioPortal (hotspots) | 3 | 3 | 6 | 3 | 44,284 <sup>(b)</sup> | 4.0 |

Figure 5 shows the distribution of these different types of variants across each protein sequence. We observe that the vast majority of variants in ATM, BRCA1 and BRCA2 are localized in flexible and/or intrinsically disordered regions (IDRs), outside Pfam domains (Figure 5, panel A). The IDRs are common in protein interaction interfaces. In fact, the NetDDR interaction network revealed a high number of interactions involving these specific DDRs: ATM (AvrNodDeg= 228), BRCA1 (AvrNodDeg= 299), BRCA2 (AvrNodDeg= 94), and MUTYH (AvrNodDeg= 17). Interestingly, the high number of interactions in ATM and BRCA1, in comparison with the moderate number in BRCA2 and MUTYH, coincide with a higher predicted consensus disorder content in ATM and BRCA1 (16.4% and 82.3%, respectively) than in BRCA2 and MUTYH (3.2% and 7.5%, respectively). The percentages of disordered regions are as given in the MobiDB database (https://mobidb.bio.unipd.it).

**Figure 5:**
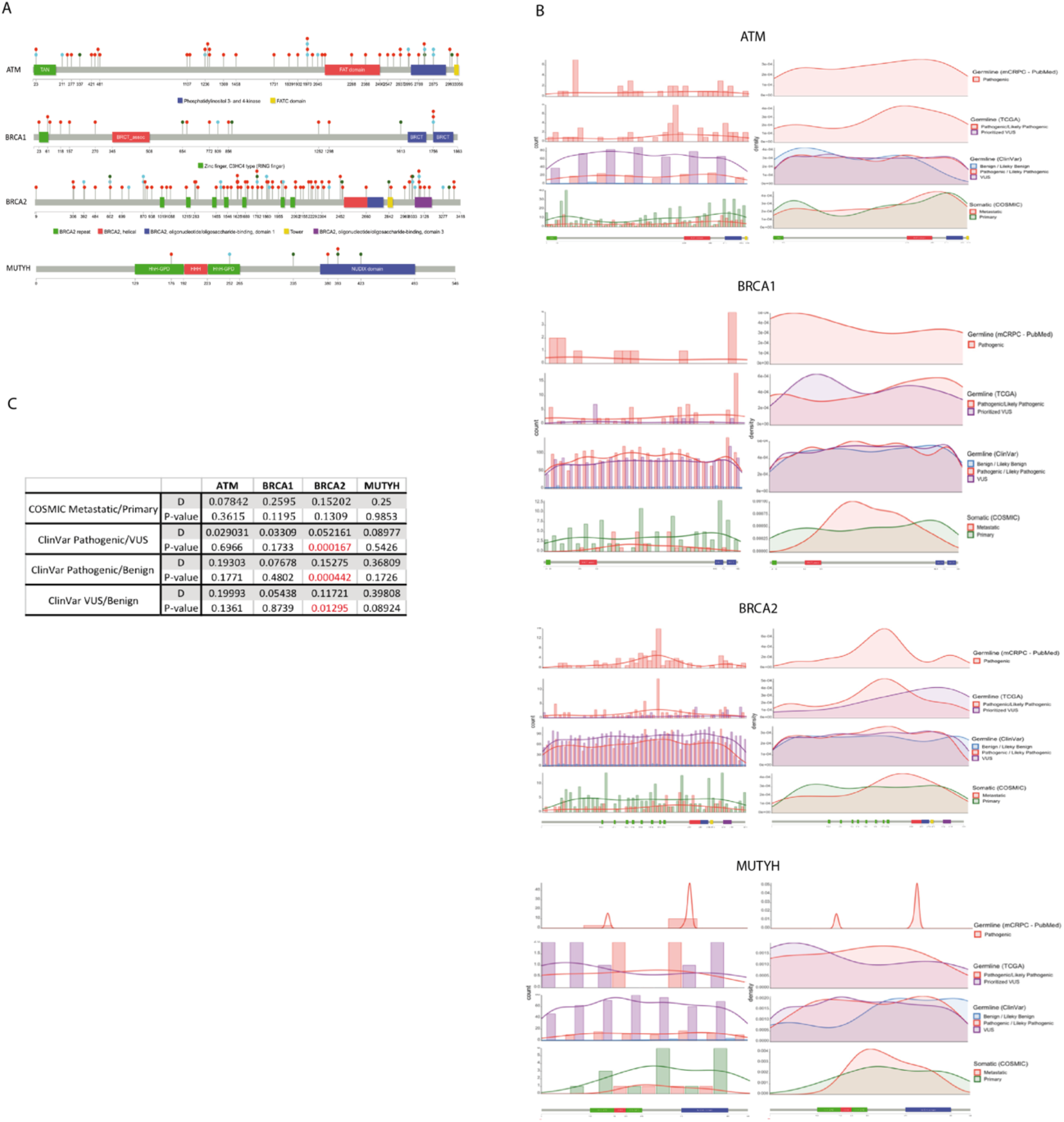
Co-localization of PCa germline and somatic variants in ATM, BRCA1, BRCA2 and MUTYH. Panel A: Location of pathogenic germline variants in different cohorts (red), recurrent germline and somatic variants in PCa (cyan), hotspot positions in different tumour types (green), and domains are shown by Lollipop structures. Panel B: Histogram and density plots of germline and somatic variants. Panel C: statistical significance between distributions.

Moreover, we observed a few cases where PCa germline and recurrent somatic variants either co-localize at the same amino acid position, or are close neighbours (e.g. red and cyan dots in Figure 5, panel A). Based on this finding, we expanded the analysis and compared the distribution of germline and somatic variants from different datasets as shown in Figure 5, panel B. Germline and somatic variants are distributed along the full length of the protein sequence, although the shape of histograms and density plots is not symmetrical and shows different peaks, which indicate accumulation of variants in defined regions of the protein. Figure 5, panel C, shows significant differences (P-value smaller than 0.05) between the pathogenic germline variants versus their benign and VUS counterparts in BRCA2. The same significant tendency was observed in BRCA2 for the distribution of VUS versus benign, both in the ClinVar dataset. No significant differences were observed in the distribution of somatic variants between primary and metastatic tumours.

In ATM, the pathogenic germline variants from different PCa cohorts, TCGA, and ClinVar datasets show a tendency to accumulate in the interaction region with the NF-κ B essential modulator IKBKG (NEMO) (a.a. 1960-2565; IntAct accessions: EBI-495465 and EBI-81279) that overlaps a flexible region flanking the FATC domain (Pfam code: PF02260). The density of benign variants in this interaction region is significantly different from pathogenic and VUS. Somatic variants from COSMIC in primary and metastatic tumours also tend to accumulate in this interaction region. A different scenario is observed in BRCA1 and BRCA2. In BRCA1, the pathogenic germline variants from ClinVar show a uniform distribution, but in different cohorts of PCa they accumulate over a flexible region connecting Zinc finger RING type (zf-C3HC4; Pfam code: PF00097) and the serine-rich associated with BRCT (BRCT_assoc; Pfam code: PF12820) N-terminal domains. By contrast, pathogenic variants from TCGA accumulate in the C-terminus, which is the interaction region with FANCJ (IntAct accessions: EBI-3509650 and EBI-349905). In BRCA2, pathogenic variants in different PCa cohorts, TCGA, and ClinVar datasets show a peak over the central flexible region including BRCA2 repeats (Pfam code: PF00634) that constitute the interaction region with RAD51 (IntAct accessions: EBI-79792 and EBI-15557721). Somatic variants in metastatic tumours accumulate in the linker and flexible region between central BRCA2 repeats and the C-terminus of the protein. These findings reinforce the idea of a synergistic effect between germline and somatic variants, and that somatic events tend to accumulate in protein interaction regions such as IDRs. Also, according to the mutational data available, for some DDR members it is possible to identify protein regions that accumulate pathogenic variants versus benign and VUS. Remarkably, disrupting or impairing these protein interactions is likely to have a marked impact on their function because these cancer-predisposition genes are highly connected and central in the NetDDR interaction network.

We also studied the co-localization of variants that are discontinuous along the sequence but proximal in the protein 3D structure. We identified different clusters containing at least three individual positions within spheres with a 15-30Å diameter that accommodate germline and somatic variants (Figure 6 and Table 5). Variants included in the same 3D-cluster can be considered part of a continuum of cancer-promoting variants, each with a relatively small but additive effect.

**Figure 6:**
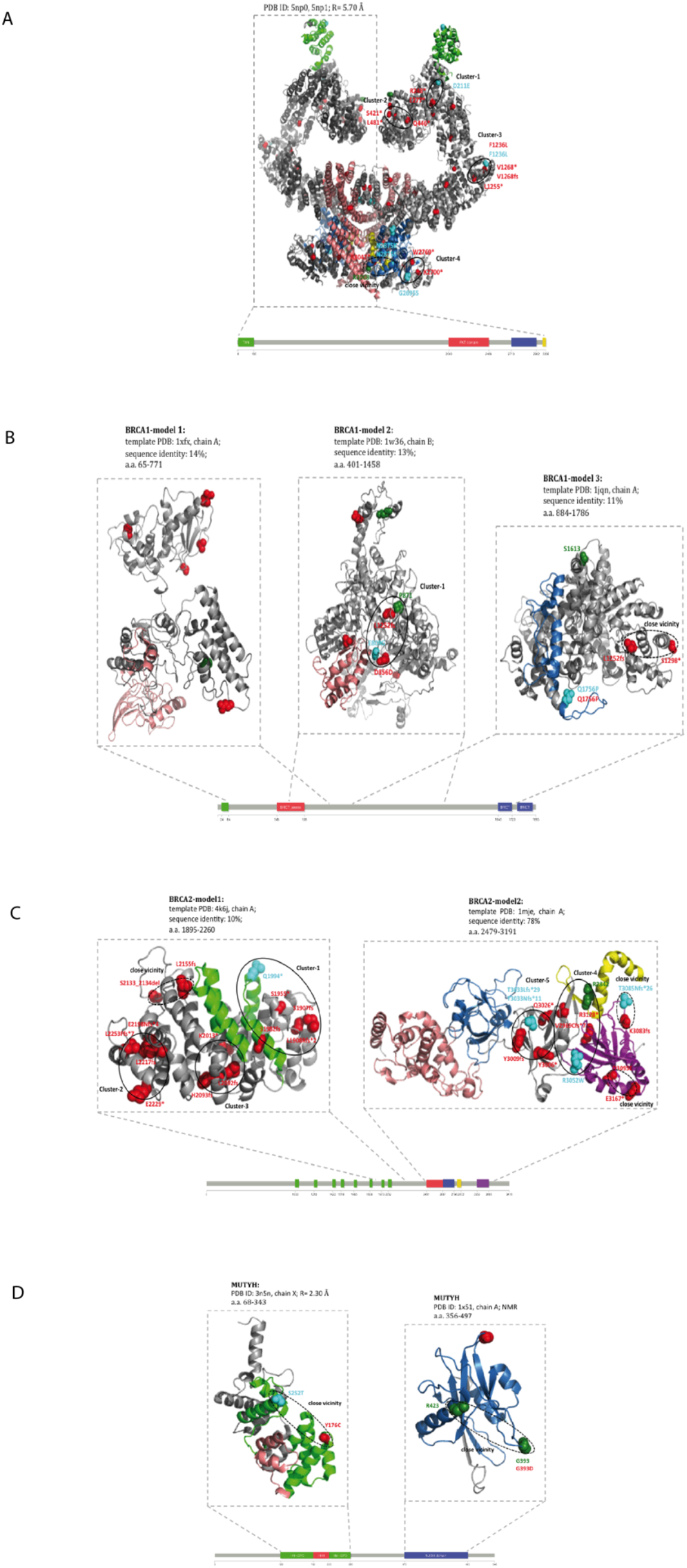
Mapping of germline and somatic variants onto protein 3D structures. The spatial 3D clusters in ATM (panel A), BRCA1 (panel B), BRCA2 (panel C), and MUTYH (panel D) are highlighted. Pathogenic germline variants are represented in red, recurrent somatic and germline in PCa in cyan, and hotspot positions in different tumour types in green. The Pfam domain colour is the same as in the Lolliplot diagrams in Figure 5.

The pathogenic germline variants in PCa, recurrent germline and somatic variants, and hotspot positions in different tumour types were mapped onto ATM, BRCA1, BRCA2 and MUTYH available structures (Figure 6) in order to find 3D-clusters of variants. According to the low-resolution Cryo-EM structures (PDB ID: 5np0, 5np1 at 5.70 Å resolution; a.a. 1-3056) we identified four 3D-clusters listed in Table 5. Other groups of variants outside 3D-clusters are positioned close to each other but with less significant P-values. As an example, the PCa pathogenic germline variant p.R3047*, recurrent somatic variants p.N2875K and p.N2875S, and the hotspot position R3008 in different tumour types are within a sphere of diameter = 25.6 Å and P-value = 0.1 (Figure 6, panel A). In the BRCA1 protein model (Modbase: a.a. 401-1458; template PDB 1w36, chain B; sequence identity 13%) we found only one 3D-cluster as listed in Table 5. Moreover, the pathogenic germline variant p.S1298* is also in close proximity to p.L1252fs but using a different 3D model (Modbase: a.a. 884-1786; template PDB 1jqn, chain A; sequence identity 11%) (diameter = 23.3 Å and P-value = 0.07) (Figure 6, panel B).

In the BRCA2 protein, we identified five 3D-clusters (see Table 5) in two 3D models from ModBase (a.a. 1895-2260; template PDB 4k6j, chain A; sequence identity 10%, and a.a. 2479-3191; template PDB 1mje, chain A; sequence identity 78%). Other variants in BRCA2 are also in close proximity: pathogenic germline p.2133_2134del and p.L2155fs (diameter = 15.5 Å and P-value = 1.0×10^−4^); recurrent somatic p.T3085Nfs*26 and pathogenic germline p.K3083fs (diameter = 6.5 Å and P-value = 1.0×10^−4^); and pathogenic germline p.D3095E, p.E3167* (diameter = 12.9657 Å and P-value = 1.0×10^−4^) (Figure 6, panel C). In the MUTYH protein, no 3D-clusters of somatic and germline variants were identified. However, according to the MUTYH crystal structure (PDB ID: 3n5n at 2.30 Å resolution; a.a. 76-362) and the solution NMR structure (PDB ID: 1×51; a.a. 356-497), pairs of variants are located in the same spatial region (Figure 6, panel D).

In the MUTYH crystal structure (PDB ID: 3n5n at 2.30 Å resolution; a.a. 76-362) the pathogenic germline variant p.Y176C is located in the same spatial region as recurrent somatic variant p.S252T in PCa (diameter = 26.4 Å and P-value = 0.7). The solution NMR structure (PDB ID: 1×51; a.a. 356-497) indicated that germline and silent variant p.G393D is located in the same surface region as hotspot positions in different tumour types G393 and R423 (diameter = 24.5 Å and P-value = 0.5) (see Figure 6, panel D). These findings suggest that the accumulation of variants in these spatial regions impairs protein interaction interfaces, and hence the biological function of the protein. The co-localization and 3D-clustering of germline and somatic variants onto the protein 3D-structure have also been applied by other authors to link rare predisposition variants to functional consequence (Y. J. Huang et al., 2018).

## Discussion

DNA repair pathways protect cells against genomic damage; disruption of these pathways can contribute to the development of cancer. In this study we show that an integrative structural analysis of affected regions in the DDR protein-coding genes can help identify susceptibility to tumour development. Based on a combined analysis of the NetDDR interaction network and the curated list of germline and somatic variants mapped onto 1,411 nodes, we first observed that the percentage of DDR and non-DDR hits with annotations in COSMIC are very similar, while in ClinVar the percentage of DDR-hits with annotations is 2-fold higher than in non-DDR. This first observation suggested some bias in the ClinVar annotations towards the most studied genes reported in the literature, which may bias research findings or limit generalizability of the results. A second observation from this analysis was the importance of the highly connected DDR genes associated with cancer-predisposition: *ATM*, *BRCA1*, *BRCA2*, *MLH1*, *MSH2*, and *MSH6*. These DDR genes are among the most affected in the COSMIC and ClinVar datasets and their encoded proteins are central in the interaction network.

Proteins in the NetDDR network range in length from 44 to 7,968 amino acid residues, therefore we analysed the accumulation of germline and somatic variants as a function of protein length. We found that DDR hits accumulate a statistically higher number of germline variants per protein length than non-DDR (P-value = 5.0×10^−4^).

Indeed, accumulation of variants is not uniform along the complete list of DDR proteins, showing a high number of germline variants accumulated in a few DDR hits. These DDR hits (BRCA1, BRCA2, MSH2, MLH1, and MSH6) coincide with highly connected and central proteins in the NetDDR interaction network. On the other hand, numbers of somatic variants per protein length are similar in the DDR and non-DDR groups (P-value = 0.36). However, genes *ATM*, *ATRX*, *CHEK2*, *ERCC2*, *MLH1*, *MSH6*, and *SMARCA4* accumulate somatic variants above the 75th percentile of the distribution.

Our analysis of germline and somatic variants in the different pathways where DDRs are involved showed that the Homology Recombination, Fanconi Anaemia, and Mismatch Repair pathways are the most affected by both types of mutations, whereas the Nucleotide Excision Repair pathway appears to be more affected by somatic variants in primary tumours than by germline pathogenic variants. The latter finding is in agreement with the results of other authors in that it shows an increased contribution of a somatic mutational pattern (Jager et al., 2019).

Relatively few articles have investigated the structural and co-localization relationships between germline and somatic variants, and none have focused on a specific protein family (Sivley et al., 2018). Hence, in this article, we analysed different structural features: Pfam domains, 3D protein interfaces, protein flexible and/or intrinsically disordered regions (IDRs), and 3D clustering of variants. Interestingly, we discovered that 47.6% of somatic variants in metastatic tumours occur within Pfam domains, which is nearly double the value of 25% for germline variants. The percentage of somatic variants in metastatic tumours affecting 3D protein interfaces (36.4%) is 3-fold higher than the value of 11% to 13% in germline variants. Furthermore, it appears that accumulation of both germline and somatic variants within Pfam domains and 3D protein interfaces results in a combined effect that damages protein function.

In this article, as use cases in the DDR study, we investigated *ATM*, *BRCA1, BRCA2* and *MUTYH* genes previously characterized in the mCRPCa PROREPAIR-B cohort (Castro et al., 2019). We analysed pathogenic germline variants identified in different cohorts of advanced PCa reported in the literature, recurrent somatic and germline variants in PCa as well as hotspot positions in different tumour types collected from cBioPortal and the TCGA-PanCancer study of pathogenic germline variants in 10,389 adult cancers (K. lin Huang et al., 2018). Using this large dataset, we observed that the vast majority of variants in ATM, BRCA1 and BRCA2 are located in flexible and/or intrinsically disordered regions (IDRs), which are common in protein interaction interfaces. Moreover, it is possible to identify protein regions where germline pathogenic variants accumulate more than benign and VUS variants. These results together reinforce the hypothesis that there is a synergistic effect between germline and somatic variants affecting protein function and interactions.

It is worth noting that a recent study into aggressive PCa, not limited to DDR genes, proposed 90 functionally related genes containing both germline and somatic mutations, and also identified biological pathways that are enriched for germline and somatic mutations – including *P53*, *STAT3*, *NKX3-1*, *KLK3*, and Androgen receptor signalling (Mamidi et al., 2019). The analysis used germline variants from genome-wide association studies (GWAS) and somatic variants identified in a cohort of 305 patients downloaded from the Cancer Genome Atlas (TCGA). Interestingly, these authors claim that genes containing germline variation did not have a high frequency of somatic mutations, which is opposite to the findings we presented here for the DDR genes. Having said that, only *ATM* is shared between ours and their analyses.

Co-localization and 3D-clustering of germline and somatic variants on the protein 3D-structure have previously been used to link rare predisposition variants to functional consequences (Y. J. Huang et al., 2018). Here we identified different clusters of variants within spheres with a 15-30Å diameter that accommodate germline and somatic variants in ATM, BRCA1 and BRCA2, and propose that variants in the same 3D-cluster are part of a continuum of cancer-promoting changes, each with a relatively small but additive effect. The integrative structural analysis discussed in this article provides a comprehensive characterisation of affected regions in DDRs and can help in the understanding of an individual’s susceptibility to tumour development.

Overall, our findings indicated a synergistic effect between germline and somatic variants affecting protein domains and interactions in the DDR family genes. In particular, Pfam domains and protein interactions interfaces are more likely to be affected by somatic variants than pathogenic germline or VUS, suggesting that the emergence of a second somatic “hit” damages the protein’s function. On the other hand, we documented 3D clusters of pathogenic germline, recurrent somatic variants from primary and metastatic tumours, and hotspots positions in ATM and BRCA2. Proper structural characterization of germline and somatic variants is needed to better stratify cancer patients with affected driver genes.

## Methods

### Protein-Protein Interaction network of DDRs

The aggregated protein-protein interaction (PPI) network for the 276 DDRs (DDR-hits) was constructed based on data from the BioGRID (Oughtred et al., 2019), InWeb_IM (Li et al., 2017) and OmniPath (Türei et al., 2016) databases. These databases were searched using the Metascape tool (Zhou et al., 2019). The aggregated PPI-Network comprised 1,466 nodes (276 DDR-hits and 1,191 non-DDR-hits) and 38,494 edges. Only 268 of the 276 DDR proteins have at least one interaction among themselves, and 229 of these 268 were considered to have enriched interactions or be “over-connected”. According to Metascape, an “over-connected” protein means that it has more than two interactions with other DDR-hits and possesses an over-connection p-value <0.01 (Zhou et al., 2019). In brief, for each connected network component, Metascape iteratively applies the MCODE algorithm to identify densely connected complexes; therefore, less-connected proteins in the initial aggregated PPI-Network are excluded from subsequent analyses. This way we get the more relevant DDR-hits, and at the same time, we reduce false positives in protein interaction data and prevent unnecessary expansion and establishment of random associations. The final and more connected network, hereafter referred to as the ‘NetDDR’, contains 1,411 nodes (229 DDR-hits and 1,182 non-DDR-hits) and 36,522 edges. The NetworkAnalyzer plugin (http://apps.cytoscape.org/apps/networkanalyzer), was used to compute topology parameters for each node in the NetDDR undirected network (Supplementary Table 1).

### Functional enrichment and pathway analysis

A pathway and biological function enrichment analysis was done for the 268 over-connected proteins using the Metascape web tool (https://metascape.org). For the enrichment analysis we selected diverse ontologies such as: KEGG Functional Sets, Pathways, and Structural Complexes; GO Molecular Functions, Biological Processes, and Cellular Components, Oncogenic Signatures, Reactome Gene Sets, Canonical Pathways, BioCarta Gene Sets, CORUM and DisGeNET (Supplementary Figure 1).

### Detection of densely connected regions in the NetDDR network

We used the MCODE algorithm, implemented in Metascape, to identify the highly interconnected proteins (or clusters) in the NetDDR network. The Metascape implementation of MCODE, can also make a GO enrichment analysis for each cluster identified by MCODE to assign “meanings” to the subnetwork or cluster components (Supplementary Figure 2), where the top three best p-value terms were retained.

### A curated dataset of germline and somatic variants for the DDR genes and protein interactors in the NetDDR network

A curated dataset of somatic and germline variants was collected for the DDR genes involved in the NetDDR network, and also for their protein interactors (229 DDR-hits and 1,182 non-DDR-hits) from the COSMIC (https://cancer.sanger.ac.uk/cosmic) and ClinVar (https://www.ncbi.nlm.nih.gov/clinvar/) databases. Careful manual curation was required to compile a high-quality dataset of variants and to end up with an accurate set in each category (i.e. confirmed somatic, primary tumour or metastasis, confirmed germline, pathogenic, benign, unknown significance). After analysing the annotations, the selected variants, together with variants identified in different cohorts collected from the literature (https://pubmed.ncbi.nlm.nih.gov/) and TCGA (https://www.cancer.gov/tcga), were used to study the distributions across the protein sequences, Pfam domains, and protein 3D structures (see Supplementary File 1).

*Somatic variants in COSMIC:* cancer-associated somatic variants were selected from COSMIC v90 (Forbes et al., 2015). This includes over 32,000 genomes, consisting of peer reviewed large-scale genome screening, and variants reported in The Cancer Genome Atlas (TCGA) and International Cancer Genome Consortium (ICGC). Only confirmed somatic variants were used when extracting variants associated with the selected DDR genes. Variants with undefined annotations (unclear syntax), as well as variants in non-coding regions, and synonymous variants were excluded. In order to focus on putative driver variants, only the variants identified in two or more tumour samples were considered for further analysis (Shihab et al., 2013). The pipeline described in this section is depicted in Supplementary Figure 3.

*Somatic and germline variants collected from the literature:* pathogenic germline variants identified in *ATM*, *BRCA1*, *BRCA2*, and *MUTYH* in the PROREPAIR-B Spanish cohorts and other mCRPCa published cohorts (see Table 1 and Supplementary File 1), were collected for the analysis. In addition, a non-redundant dataset from 176 different studies containing 46,566 non-overlapping samples from 44,284 patients was explored using cBioPortal (https://www.cbioportal.org) (Cerami et al., 2012; Gao et al., 2013). Recurrent somatic and germline variants in PCa (n = 1,556) and hotspot positions in different tumour types (n = 44,284) in these genes are documented in Supplementary File 1.

*Germline variants in ClinVar*: variants clearly classified as germline, according to *ClinVar* (Aug 2019), were annotated using VEP (McLaren et al., 2016) to check their genomic location and to include only the variants in protein coding regions. After discarding the variants with undefined or unclear annotation, we categorized the remaining variants into three different groups according to the Clinical Significance annotations provided by ClinVar. The groups were “pathogenic/likely pathogenic”, “benign/likely benign”, and “variants with unknown significance (VUS)”. For more details, see Supplementary Figure 4.

*Germline variants from TCGA*: germline variants classified as pathogenic or prioritized VUS were collected from TCGA (Knijnenburg et al., 2018). We used *ATM*, *BRCA1*, *BRCA2*, and *MUTYH* as a proxy for the TCGA dataset (see Table 1 and Supplementary File 1).

### Structural modelling of proteins

Before mapping the germline and somatic variants onto the 3D-structures of PCa predisposition genes *ATM, BRCA1*, *BRCA2*, and *MUTYH*, we reviewed the RCSB PDB database (https://www.rcsb.org), which collects experimentally determined 3D-structures, and the ModBase repository (http://modbase.compbio.ucsf.edu) which generates homology-built protein 3D models. In the latter, the sequence identity between target proteins and those whose crystal structures were used as a template, ranged from 10% to 100%.

Human ATM (3,056 a.a. UniProt: Q13315), BRCA1 (1,863 a.a. UniProt: P38398), BRCA2 (3,418 a.a. UniProt: P51587) and MUTYH (546 a.a. Unirpot: Q9UIF7) are multi-domain proteins whose complete crystallographic structure has not been determined yet. However, X-ray crystal structures (PDB id: 1jnx, 1n5o, 1t15, 1t29, 3eu7, 1n0w, 3n5n, 3n5n, 1×51) and solution structures (PDB id: 6hka, 1jm7, 1oqa, 1×51, 5dpk) of different regions of these proteins are available. Furthermore, low-resolution electron microscopy models (PDB id: 5np0, 5np1) covering the complete sequence of ATM have been published. Additional data about the resolution, protein chains, and amino acid regions are shown in **Table 2**.

**Table 2:** List of the 3D structures of the key DDR proteins available from the PDB database.

|  | Region | Method | PDB id | Resolution | Chains |
| --- | --- | --- | --- | --- | --- |
| ATM | 1-3056 | Cryo-EM | 5np0 | 5.70 Å | A/B |
|  | 1-3056 | Cryo-EM | 5np1 | 5.70 Å | A |
|  | 3024-3030 | NMR | 6hka | - | A |
| BRCA1 | 1-110 | NMR | 1JM7 | - | A |
|  | 1646-1859 | X-ray | 1JNX | 2.50 Å | X |
|  | 1646-1859 | X-ray | 1N50 | 2.80 Å | X |
|  | 1755-1863 | NMR | 1OQA | - | A |
|  | 1646-1859 | X-ray | 1T15 | 1.85 Å | A |
|  | 1646-1859 | X-ray | 1T29 | 2.30 Å | A |
| BRCA2 | 1517-1551 | X-ray | 1N0W | 1.70 Å | B |
|  | 21-39 | X-ray | 3EU7 | 2.20 Å | A |
| MUTYH | 76-362 | X-ray | 3N5N | 2.30 Å | X/Y |
|  | 356-497 | NMR | 1X51 | - | A |

**Table 3:**
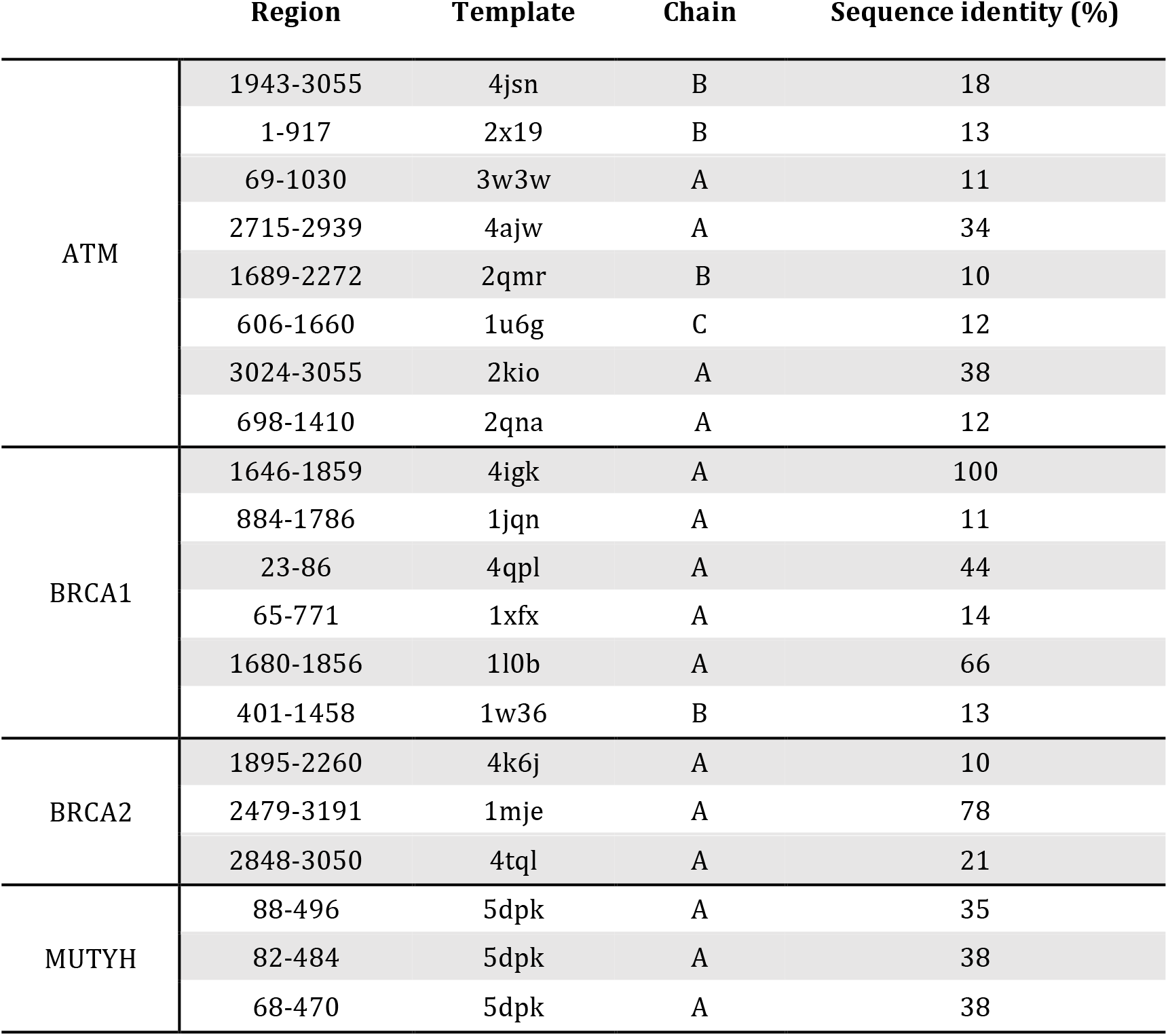
List of 3D models annotated in the ModBase database.

**Table 4:** List of Pfam domains affected by germline and somatic mutations in the DDR family.

| **Pfam ID** | **Description** | **Cellular functions of domain** |
| --- | --- | --- |
| PF05192 | MutS_III | DNA mismatch repair and recombination |
| PF05190 | MutS domain IV | Involved in mismatch repair |
| PF00488 | MutS domain V | Involved in mismatch repair and DNA binding |
| PF00634 | BRCA2 repeat | Involved in the repair of chromosomal damage with an important role in the error-free repair of DNA double strand breaks. Breast cancer susceptibility protein. |
| PF09169 | BRCA-2_helical | Binds DSS1 protein |
| PF09103 | BRCA-2_OB1 (BRCA2 oligonucleotide/oligosaccharide-binding domain 1) | Binds DSS1 protein |
| PF09121 | Tower domain | Essential for appropriate binding of BRCA2 to DNA. Has an important role in the tumour suppressor function of BRCA2 |
| PF09104 | BRCA-2_OB3 (BRCA2, oligonucleotide/oligosaccharide-binding domain 3) | Binds DSS1 protein |
| PF10409 | PTEN-C2 | Plays a central role in membrane binding, productively positions the catalytic part of the protein onto the membrane |
| PF00870 | P53 | DNA-binding domain |
| PF07710 | P53 tetramer |  |
| PF00097 | zf-C3HC4 (Zinc finger RING type) | Zinc fingers bind DNA, RNA, protein and/or lipid substrates. RING fingers play a key role in the ubiquitination pathway |
| PF12820 | BRCT_assoc (serine-rich associated with BRCT) | No specified function |
| PF00533 | BRCT (BRCA1 C-terminus) | Involved in cell cycle checkpoint functions responsive to DNA damage. A marker of breast cancer susceptibility |
| PF11640 | TAN (Telomere-length maintenance and DNA damage repair) | Regulates responses to DNA double-strand breaks (EC:2.7.11.1) |
| PF02259 | FAT (A novel domain in PIK-related proteins; i.e. *FRAP*, *ATM*, and *TRRAP*) | Still need to be elucidated experimentally. Usually involved in protein-protein interactions |
| PF00454 | PI3_PI4_kinase (Phosphatidylinositol 3- and 4-kinase) | Involved in cell growth, proliferation, differentiation, motility, survival and intracellular trafficking, which in turn are involved in cancer |
| PF02260 | FATC (A novel motif at the extreme C-terminus of PIK-related proteins) | Still need to be elucidated experimentally. Usually involved in protein-protein interactions |
| PF00730 | HhH-GPD (HHH-GPD superfamily base excision DNA repair protein) | Involved in DNA repair functions (i.e. endonuclease III, DNA glycosylase, and methyl-CPG binding protein) |
| PF00633 | HHH (Helix-hairpin-helix motif) | DNA-binding domain |
| PF14815 | NUDIX_4 (NUDIX domain) | A/G-specific adenine glycosylases. Involved in DNA base-pair mismatch repair |

**Table 5:**
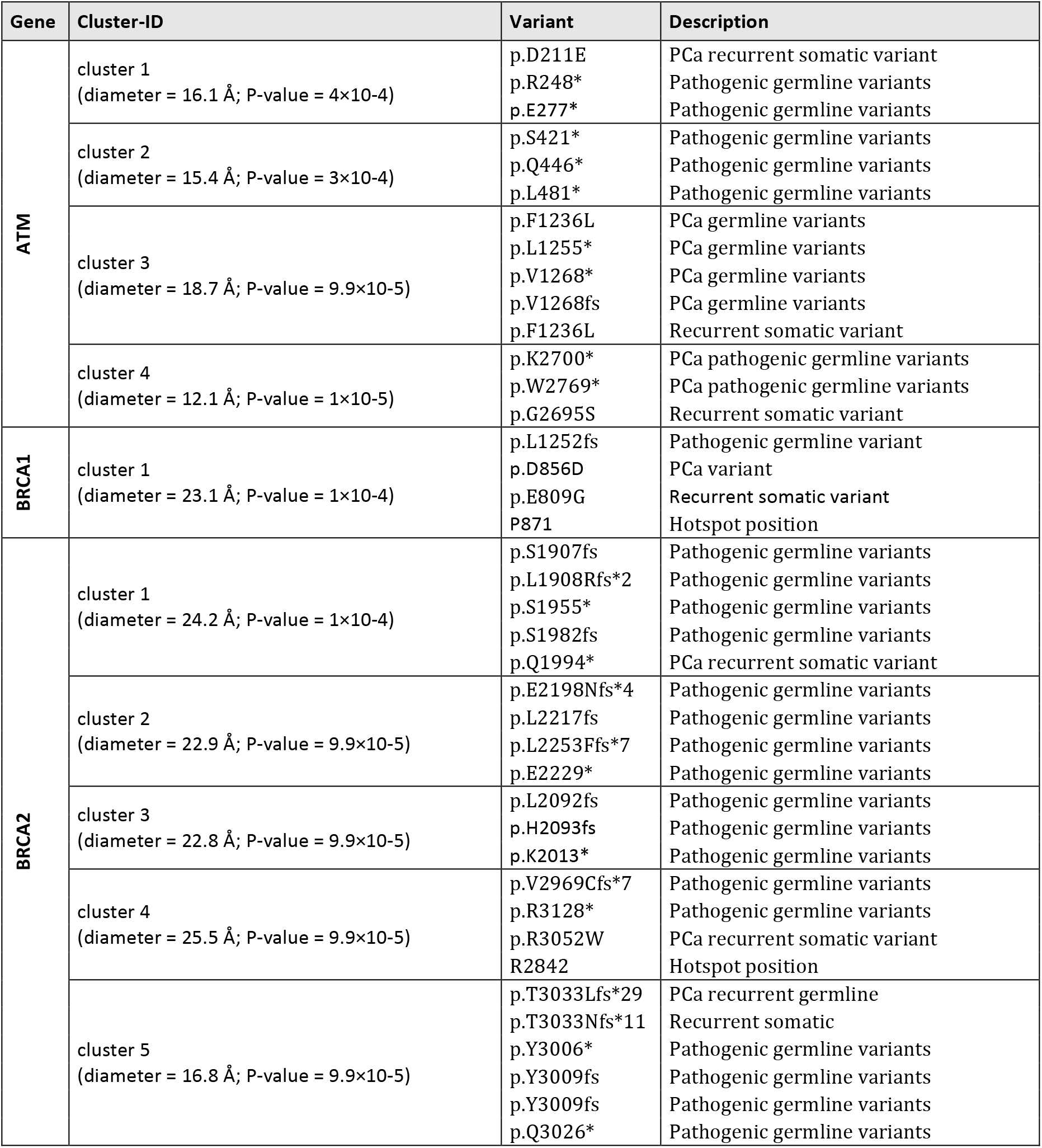
List of 3D clusters identified in the protein structure of ATM, BRCA1, BRCA2 and MUTYH.

| Gene | Cluster-ID | Variant | Description |
| --- | --- | --- | --- |
| ATM | cluster 1<br>(diameter = 16.1 Å; P-value = $4 \times 10^{-4}$) | p.D211E<br>p.R248*<br>p.E277* | PCa recurrent somatic variant<br>Pathogenic germline variants<br>Pathogenic germline variants |
| | cluster 2<br>(diameter = 15.4 Å; P-value = $3 \times 10^{-4}$) | p.S421*<br>p.Q446*<br>p.L481* | Pathogenic germline variants<br>Pathogenic germline variants<br>Pathogenic germline variants |
| | cluster 3<br>(diameter = 18.7 Å; P-value = $9.9 \times 10^{-5}$) | p.F1236L<br>p.L1255*<br>p.V1268*<br>p.V1268fs<br>p.F1236L | PCa germline variants<br>PCa germline variants<br>PCa germline variants<br>PCa germline variants<br>Recurrent somatic variant |
| | cluster 4<br>(diameter = 12.1 Å; P-value = $1 \times 10^{-5}$) | p.K2700*<br>p.W2769*<br>p.G2695S | PCa pathogenic germline variants<br>PCa pathogenic germline variants<br>Recurrent somatic variant |
| BRCA1 | cluster 1<br>(diameter = 23.1 Å; P-value = $1 \times 10^{-4}$) | p.L1252fs<br>p.D856D<br>p.E809G<br>P871 | Pathogenic germline variant<br>PCa variant<br>Recurrent somatic variant<br>Hotspot position |
| BRCA2 | cluster 1<br>(diameter = 24.2 Å; P-value = $1 \times 10^{-4}$) | p.S1907fs<br>p.L1908Rfs*2<br>p.S1955*<br>p.S1982fs<br>p.Q1994* | Pathogenic germline variants<br>Pathogenic germline variants<br>Pathogenic germline variants<br>Pathogenic germline variants<br>PCa recurrent somatic variant |
| | cluster 2<br>(diameter = 22.9 Å; P-value = $9.9 \times 10^{-5}$) | p.E2198Nfs*4<br>p.L2217fs<br>p.L2253Ffs*7<br>p.E2229* | Pathogenic germline variants<br>Pathogenic germline variants<br>Pathogenic germline variants<br>Pathogenic germline variants |
| | cluster 3<br>(diameter = 22.8 Å; P-value = $9.9 \times 10^{-5}$) | p.L2092fs<br>p.H2093fs<br>p.K2013* | Pathogenic germline variants<br>Pathogenic germline variants<br>Pathogenic germline variants |
| | cluster 4<br>(diameter = 25.5 Å; P-value = $9.9 \times 10^{-5}$) | p.V2969Cfs*7<br>p.R3128*<br>p.R3052W<br>R2842 | Pathogenic germline variants<br>Pathogenic germline variants<br>PCa recurrent somatic variant<br>Hotspot position |
| | cluster 5<br>(diameter = 16.8 Å; P-value = $9.9 \times 10^{-5}$) | p.T3033Lfs*29<br>p.T3033Nfs*11<br>p.Y3006*<br>p.Y3009fs<br>p.Y3009fs<br>p.Q3026* | PCa recurrent germline<br>Recurrent somatic<br>Pathogenic germline variants<br>Pathogenic germline variants<br>Pathogenic germline variants<br>Pathogenic germline variants |

### Identification of affected protein functional domains, key residues, protein interaction interfaces, and 3D-clusters of germline and somatic variants

Protein domain annotations and flexible, or intrinsically disordered regions (IDRs), were extracted from the Pfam (https://pfam.xfam.org) and MobiDB (https://mobidb.bio.unipd.it) databases, respectively. Information on key functional residues (i.e. catalytic, ligand-binding, posttranslational modification) was collected from UniProt (https://www.uniprot.org) using the Structure-PPi system (http://rbbt.bsc.es/Structure/Structure) (Vázquez et al., 2015). Residues in protein interaction interfaces were identified via the Structure workflow implemented in the PCAWG-Scout online browser (http://pcawgscout.bsc.es) (Campbell et al., 2020). We also used the mutation3D program (http://mutation3d.org) (Meyer et al., 2016) to identify clusters of germline and somatic variants in the PDB and ModBase models. We ran mutation3D using the command recommended by its authors (./mutation3d <pdb_file><amino acid substitutions><CL-distance><protein length><number of bootstrapping iterations>) and parameters were set as follow: diameter or CL-distance = 30.0 Å, and number of bootstrapping iterations = 10000.

## Acknowledgments

This research was funded in part by the Spanish Ministry of Science and Innovation, co-funded by FEDER funds [Severo Ochoa SVP-2014-068895 contract (L.M-P)] and EMBO Short-Term Fellowship [grant number 8374 (L.M-P)]. We also thank all members of the Thornton research group (EMBL-EBI) and the Prostate Cancer Clinical Unit (CNIO) for early feedback on our manuscript.

## Author contributions statement

L.M-P, J.M-T and R.A-L design the study. L.M-P, R.A-L, and T.P carried out the analysis. L.M-P, R.A-L, and T.P drafting of the manuscript with input from all authors. J.M-T conceptualized experiments, supervised the work, and secured funding. All authors reviewed the manuscript.

## Additional Data

Supplementary Figure 1. **Functions and pathways enrichment analysis provided by Metascape**. The Network represents the top 20 clusters with their representative enriched terms—one per cluster. (A) Each node indicates an enriched term and is coloured by its cluster identity (i.e., nodes of the same colour belong to the same cluster). Node size is proportional to the number of input genes fall into that term. Terms with a similarity score > 0.3 are linked by an edge (the thickness of the edge represents the similarity score). One term from each cluster is selected to have its term description shown as label. (B) The same network but its nodes are colored by p-value, as shown in the legend. The dark the color, the more statistically significant the node is. (C) List of the top 20 statistically enriched terms. The terms can be GO/KEGG terms, canonical pathways, hall mark gene sets, etc.

Supplementary Figure 2. **MCODE analysis of the NetDDR network as provided by Metascape**. Densely connected proteins (MCODE clusters) are highlighted. Each network is assigned a unique color. GO enrichment analysis was applied to each cluster to assign “meanings” to the network component, where top three best p-value terms were retained.

Supplementary Figure 3. **Flow chart for the classification of somatic variants extracted from COSMIC database**. DDR-hits and non-DDR-hits are according to the NetDDR network.

Supplementary Figure 4. **Flow chart for the classification of germline variants extracted from ClinVar database**. DDR-hits and non-DDR-hits are according to the NetDDR network.

Supplementary Figure 5. **Summary of germline and somatic variants extracted from ClinVar and COSMIC databases**. (A) Barplot of the selected germline variants identified in DDR genes and extracted from the ClinVar database. (B) Subset of pathogenic germline variants (only). (C) Barplot of the selected somatic variants identified in DDR genes and extracted from the COSMIC database. For a better visualization of the barplot, we exclude TP53, IDH1 and PTEN that accumulate a large number of variants.

Supplementary Table 1. **Topological parameters of the NetDDR network**.

Supplementary Table 2. **Distribution of germline and somatic variants in Pfam domains and protein interfaces**

Supplementary_File1 (MS Excel document). List of germline variants extracted from PubMed (pathogenic), TCGA (pathogenic and VUS), ClinVar (Pathogenic, Benign and VUS), and somatic variants from COSMIC (metastatic and primary) and cBioPortal (germline and recurrent somatic variants).

Supplementary_File2 (MS Excel document). Functional annotations of germline and somatic variants using the Structure-PPI system.

## References

Andrés-León, E., Cases, I., Arcas, A., & Rojas, A. M. (2016). DDRprot: A database of DNA damage response-related proteins. Database: The Journal of Biological Databases and Curation, 2016. https://doi.org/10.1093/database/baw123

Annala, M., Vandekerkhove, G., Khalaf, D., Taavitsainen, S., Beja, K., Warner, E. W., Sunderland, K., Kollmannsberger, C., Eigl, B. J., Finch, D., Oja, C. D., Vergidis, J., Zulfiqar, M., Azad, A. A., Nykter, M., Gleave, M. E., Wyatt, A. W., & Chi, K. N. (2018). Circulating Tumor DNA Genomics Correlate with Resistance to Abiraterone and Enzalutamide in Prostate Cancer. Cancer Discov, 8(4), 444–457. https://doi.org/10.1158/2159-8290.Cd-17-0937

Antonarakis, E. S., Lu, C., Luber, B., Liang, C., Wang, H., Chen, Y., Silberstein, J. L., Piana, D., Lai, Z., Chen, Y., Isaacs, W. B., & Luo, J. (2018). Germline DNA-repair Gene Mutations and Outcomes in Men with Metastatic Castration-resistant Prostate Cancer Receiving First-line Abiraterone and Enzalutamide. Eur Urol, 74(2), 218–225. https://doi.org/10.1016/j.eururo.2018.01.035

Campbell, P. J., Getz, G., Korbel, J. O., Stuart, J. M., Jennings, J. L., Stein, L. D., Perry, M. D., Nahal-Bose, H. K., Ouellette, B. F. F., Li, C. H., Rheinbay, E., Nielsen, G. P., Sgroi, D. C., Wu, C.-L., Faquin, W. C., Deshpande, V., Boutros, P. C., Lazar, A. J., Hoadley, K. A., … The ICGC/TCGA Pan-Cancer Analysis of Whole Genomes Consortium. (2020). Pan-cancer analysis of whole genomes. Nature, 578(7793), 82–93. https://doi.org/10.1038/s41586-020-1969-6

Castro, E., Romero-Laorden, N., Del Pozo, A., Lozano, R., Medina, A., Puente, J., Piulats, J. M., Lorente, D., Saez, M. I., Morales-Barrera, R., Gonzalez-Billalabeitia, E., Cendon, Y., Garcia-Carbonero, I., Borrega, P., Mendez Vidal, M. J., Montesa, A., Nombela, P., Fernandez-Parra, E., Gonzalez Del Alba, A., … Olmos, D. (2019). PROREPAIR-B: A Prospective Cohort Study of the Impact of Germline DNA Repair Mutations on the Outcomes of Patients With Metastatic Castration-Resistant Prostate Cancer. J Clin Oncol, 37(6), 490–503. https://doi.org/10.1200/jco.18.00358

Cerami, E., Gao, J., Dogrusoz, U., Gross, B. E., Sumer, S. O., Aksoy, B. A., Jacobsen, A., Byrne, C. J., Heuer, M. L., Larsson, E., Antipin, Y., Reva, B., Goldberg, A. P., Sander, C., & Schultz, N. (2012). The cBio cancer genomics portal: An open platform for exploring multidimensional cancer genomics data. Cancer Discovery, 2(5), 401–404. https://doi.org/10.1158/2159-8290.CD-12-0095

D’Andrea, A. D., & Grompe, M. (2003). The Fanconi anaemia/BRCA pathway. Nature Reviews. Cancer, 3(1), 23–34. https://doi.org/10.1038/nrc970

Edwards, S. L., Brough, R., Lord, C. J., Natrajan, R., Vatcheva, R., Levine, D. A., Boyd, J., Reis-Filho, J. S., & Ashworth, A. (2008). Resistance to therapy caused by intragenic deletion in BRCA2. Nature, 451(7182), 1111–1115. https://doi.org/10.1038/nature06548

Forbes, S. A., Beare, D., Gunasekaran, P., Leung, K., Bindal, N., Boutselakis, H., Ding, M., Bamford, S., Cole, C., Ward, S., Kok, C. Y., Jia, M., De, T., Teague, J. W., Stratton, M. R., McDermott, U., & Campbell, P. J. (2015). COSMIC: exploring the world’s knowledge of somatic mutations in human cancer. Nucleic Acids Res, 43(Database issue), D805–11. https://doi.org/10.1093/nar/gku1075

Gao, J., Aksoy, B. A., Dogrusoz, U., Dresdner, G., Gross, B., Sumer, S. O., Sun, Y., Jacobsen, A., Sinha, R., Larsson, E., Cerami, E., Sander, C., & Schultz, N. (2013). Integrative analysis of complex cancer genomics and clinical profiles using the cBioPortal. Science Signaling, 6(269), pl1. https://doi.org/10.1126/scisignal.2004088

Huang, K lin, Mashl, R. J., Wu, Y., Ritter, D. I., Wang, J., Oh, C., Paczkowska, M., Reynolds, S., Wyczalkowski, M. A., Oak, N., Scott, A. D., Krassowski, M., Cherniack, A. D., Houlahan, K. E., Jayasinghe, R., Wang, L. B., Zhou, D. C., Liu, D., Cao, S., … Ding, L. (2018). Pathogenic Germline Variants in 10,389 Adult Cancers. Cell, 173(2), 355–370.e14. https://doi.org/10.1016/j.cell.2018.03.039

Huang, Y. J., Brock, K. P., Sander, C., Marks, D. S., & Montelione, G. T. (2018). A Hybrid Approach for Protein Structure Determination Combining Sparse NMR with Evolutionary Coupling Sequence Data. Advances in Experimental Medicine and Biology, 1105, 153–169. https://doi.org/10.1007/978-981-13-2200-6_10

Jager, M., Blokzijl, F., Kuijk, E., Bertl, J., Vougioukalaki, M., Janssen, R., Besselink, N., Boymans, S., Ligt, J. de, Pedersen, J. S., Hoeijmakers, J., Pothof, J., Boxtel, R. van, & Cuppen, E. (2019). Deficiency of nucleotide excision repair is associated with mutational signature observed in cancer. Genome Research, 29(7), 1067–1077. https://doi.org/10.1101/gr.246223.118

Jonsson, P., Bandlamudi, C., Cheng, M. L., Srinivasan, P., Chavan, S. S., Friedman, N. D., Rosen, E. Y., Richards, A. L., Bouvier, N., Selcuklu, S. D., Bielski, C. M., Abida, W., Mandelker, D., Birsoy, O., Zhang, L., Zehir, A., Donoghue, M. T. A., Baselga, J., Offit, K., … Taylor, B. S. (2019). Tumour lineage shapes BRCA-mediated phenotypes. Nature, 571(7766), 576–579. https://doi.org/10.1038/s41586-019-1382-1

Khurana, E., Fu, Y., Colonna, V., Mu, X. J., Kang, H. M., Lappalainen, T., Sboner, A., Lochovsky, L., Chen, J., Harmanci, A., Das, J., Abyzov, A., Balasubramanian, S., Beal, K., Chakravarty, D., Challis, D., Chen, Y., Clarke, D., Clarke, L., … Gerstein, M. (2013). Integrative annotation of variants from 1092 humans: Application to cancer genomics. Science (New York, N.Y.), 342(6154), 1235587. https://doi.org/10.1126/science.1235587

Knijnenburg, T. A., Wang, L., Zimmermann, M. T., Chambwe, N., Gao, G. F., Cherniack, A. D., Fan, H., Shen, H., Way, G. P., Greene, C. S., Liu, Y., Akbani, R., Feng, B., Donehower, L. A., Miller, C., Shen, Y., Karimi, M., Chen, H., Kim, P., … Wang, C. (2018). Genomic and Molecular Landscape of DNA Damage Repair Deficiency across The Cancer Genome Atlas. Cell Rep, 23(1), 239–254.e6. https://doi.org/10.1016/j.celrep.2018.03.076

Lans, H., Hoeijmakers, J. H. J., Vermeulen, W., & Marteijn, J. A. (2019). The DNA damage response to transcription stress. Nature Reviews Molecular Cell Biology, 20(12), 766–784. https://doi.org/10.1038/s41580-019-0169-4

Li, T., Wernersson, R., Hansen, R. B., Horn, H., Mercer, J., Slodkowicz, G., Workman, C. T., Rigina, O., Rapacki, K., Stærfeldt, H. H., Brunak, S., Jensen, T. S., & Lage, K. (2017). A scored human protein–protein interaction network to catalyze genomic interpretation. Nature Methods, 14(1), 61–64. https://doi.org/10.1038/nmeth.4083

Lu, C., Xie, M., Wendl, M. C., Wang, J., McLellan, M. D., Leiserson, M. D., Huang, K. L., Wyczalkowski, M. A., Jayasinghe, R., Banerjee, T., Ning, J., Tripathi, P., Zhang, Q., Niu, B., Ye, K., Schmidt, H. K., Fulton, R. S., McMichael, J. F., Batra, P., … Ding, L. (2015). Patterns and functional implications of rare germline variants across 12 cancer types. Nat Commun, 6, 10086–10086. https://doi.org/10.1038/ncomms10086

Mamidi, T. K. K., Wu, J., & Hicks, C. (2019). Mapping the Germline and Somatic Mutation Interaction Landscape in Indolent and Aggressive Prostate Cancers. Journal of Oncology, 2019, 4168784. https://doi.org/10.1155/2019/4168784

McLaren, W., Gil, L., Hunt, S. E., Riat, H. S., Ritchie, G. R. S., Thormann, A., Flicek, P., & Cunningham, F. (2016). The Ensembl Variant Effect Predictor. Genome Biology, 17(1), 122. https://doi.org/10.1186/s13059-016-0974-4

Meyer, M. J., Lapcevic, R., Romero, A. E., Yoon, M., Das, J., Beltrán, J. F., Mort, M., Stenson, P. D., Cooper, D. N., Paccanaro, A., & Yu, H. (2016). mutation3D: Cancer Gene Prediction Through Atomic Clustering of Coding Variants in the Structural Proteome. Human Mutation, 37(5), 447–456. https://doi.org/10.1002/humu.22963

Mijuskovic, M., Saunders, E. J., Leongamornlert, D. A., Wakerell, S., Whitmore, I., Dadaev, T., Cieza-Borrella, C., Govindasami, K., Brook, M. N., Haiman, C. A., Conti, D. V., Eeles, R. A., & Kote-Jarai, Z. (2018). Rare germline variants in DNA repair genes and the angiogenesis pathway predispose prostate cancer patients to develop metastatic disease. Br J Cancer, 119(1), 96–104. https://doi.org/10.1038/s41416-018-0141-7

Na, R., Zheng, S. L., Han, M., Yu, H., Jiang, D., Shah, S., Ewing, C. M., Zhang, L., Novakovic, K., Petkewicz, J., Gulukota, K., Helseth Jr., D. L., Quinn, M., Humphries, E., Wiley, K. E., Isaacs, S. D., Wu, Y., Liu, X., Zhang, N., … Isaacs, W. B. (2017). Germline Mutations in ATM and BRCA1/2 Distinguish Risk for Lethal and Indolent Prostate Cancer and are Associated with Early Age at Death. Eur Urol, 71(5), 740–747. https://doi.org/10.1016/j.eururo.2016.11.033

Oughtred, R., Stark, C., Breitkreutz, B.-J., Rust, J., Boucher, L., Chang, C., Kolas, N., O’Donnell, L., Leung, G., McAdam, R., Zhang, F., Dolma, S., Willems, A., Coulombe-Huntington, J., Chatr-Aryamontri, A., Dolinski, K., & Tyers, M. (2019). The BioGRID interaction database: 2019 update. Nucleic Acids Research, 47(D1), D529–D541. https://doi.org/10.1093/nar/gky1079

Pritchard, C. C., Mateo, J., Walsh, M. F., De Sarkar, N., Abida, W., Beltran, H., Garofalo, A., Gulati, R., Carreira, S., Eeles, R., Elemento, O., Rubin, M. A., Robinson, D., Lonigro, R., Hussain, M., Chinnaiyan, A., Vinson, J., Filipenko, J., Garraway, L., … Nelson, P. S. (2016). Inherited DNA-Repair Gene Mutations in Men with Metastatic Prostate Cancer. N Engl J Med, 375(5), 443–453. https://doi.org/10.1056/NEJMoa1603144

Robinson, D., Van Allen, E. M., Wu, Y. M., Schultz, N., Lonigro, R. J., Mosquera, J. M., Montgomery, B., Taplin, M. E., Pritchard, C. C., Attard, G., Beltran, H., Abida, W., Bradley, R. K., Vinson, J., Cao, X., Vats, P., Kunju, L. P., Hussain, M., Feng, F. Y., … Chinnaiyan, A. M. (2015). Integrative clinical genomics of advanced prostate cancer. Cell, 161(5), 1215–1228. https://doi.org/10.1016/j.cell.2015.05.001

Shihab, H. A., Gough, J., Cooper, D. N., Day, I. N. M., & Gaunt, T. R. (2013). Predicting the functional consequences of cancer-associated amino acid substitutions. Bioinformatics (Oxford, England), 29(12), 1504–1510. https://doi.org/10.1093/bioinformatics/btt182

Sivley, R. M., Dou, X., Meiler, J., Bush, W. S., & Capra, J. A. (2018). Comprehensive Analysis of Constraint on the Spatial Distribution of Missense Variants in Human Protein Structures. American Journal of Human Genetics, 102(3), 415–426. https://doi.org/10.1016/j.ajhg.2018.01.017

Sokol, E. S., Pavlick, D., Khiabanian, H., Frampton, G. M., Ross, J. S., Gregg, J. P., Lara, P. N., Oesterreich, S., Agarwal, N., Necchi, A., Miller, V. A., Alexander, B., Ali, S. M., Ganesan, S., & Chung, J. H. (2020). Pan-Cancer Analysis of BRCA1 and BRCA2 Genomic Alterations and Their Association With Genomic Instability as Measured by Genome-Wide Loss of Heterozygosity. JCO Precision Oncology. https://doi.org/10.1200/PO.19.00345

Toledo, R. A., & Group, N. G. S. in P. S. (2018). Inflated pathogenic variant profiles in the ClinVar database. Nat Rev Endocrinol, 14(7), 387–389. https://doi.org/10.1038/s41574-018-0034-0

Türei, D., Korcsmáros, T., & Saez-Rodriguez, J. (2016). OmniPath: Guidelines and gateway for literature-curated signaling pathway resources. Nature Methods, 13(12), 966–967. https://doi.org/10.1038/nmeth.4077

Vázquez, M., Valencia, A., & Pons, T. (2015). Structure-PPi: A module for the annotation of cancer-related single-nucleotide variants at protein–protein interfaces. Bioinformatics, 31(14), 2397–2399. https://doi.org/10.1093/bioinformatics/btv142

Zhou, Y., Zhou, B., Pache, L., Chang, M., Khodabakhshi, A. H., Tanaseichuk, O., Benner, C., & Chanda, S. K. (2019). Metascape provides a biologist-oriented resource for the analysis of systems-level datasets. Nature Communications, 10(1), 1–10. https://doi.org/10.1038/s41467-019-09234-6

