## Supplementary Tables and Figures for "A computational and structural analysis of germline and somatic variants affecting the DDR mechanism, and their impact on human diseases and prostate cancer progression"

Supplementary Table 1. **Topological parameters of the NetDDR network**

| **Parameters** | **1,411 nodes (NetDDR)** | **229 nodes (DDR)** | **1,182 nodes (non-DDR)** |
|---|---|---|---|
| AverageShortestPathLength | 2.36 | 2.2 | 2.39 |
| ClosenessCentrality | 0.43 | 0.46 | 0.42 |
| ClusteringCoefficient | 0.36 | 0.39 | 0.35 |
| Degree | 51.77 | 87.17 | 44.91 |
| Eccentricity | 3.7 | 3.52 | 3.74 |
| NeighborhoodConnectivity | 133.66 | 125.05 | 135.32 |
| Radiality | 0.73 | 0.76 | 0.72 |
| Stress | 70391.78 | 183985.76 | 48384.15 |
| TopologicalCoefficient | 0.18 | 0.13 | 0.19 |

Supplementary Table 2. **Distribution of germline and somatic variants in Pfam domains and protein interfaces**

| | | Total | No annotation | Outside Pfam | Inside Pfam | Interfaces |
|---|---|---|---|---|---|---|
| Germline (ClinVar) | Pathogenic | 10, 301 | 532 (5.2%) | 7 247 (74.2%) | 2 522 (25.8%) | 1 268 (12.9%) |
| | Benign | 1, 117 | 123 (11%) | 857 (86.2%) | 137 (13.8%) | 72 (7.2%) |
| | VUS | 28 248 | 3 737 (13.2%) | 18,375 (75%) | 6,136 (25%) | 2,788 (11.4%) |
| Somatic (COSMIC) | Metastasis | 2,030 | 490 (24.1%) | 807 (52.4%) | 733 (47.6%) | 561 (36.4%) |
| | Primary | 5,795 | 1,297 (22%) | 2,830 (62.9%) | 1,668 (37.1%) | 1,058 (23.5%) |

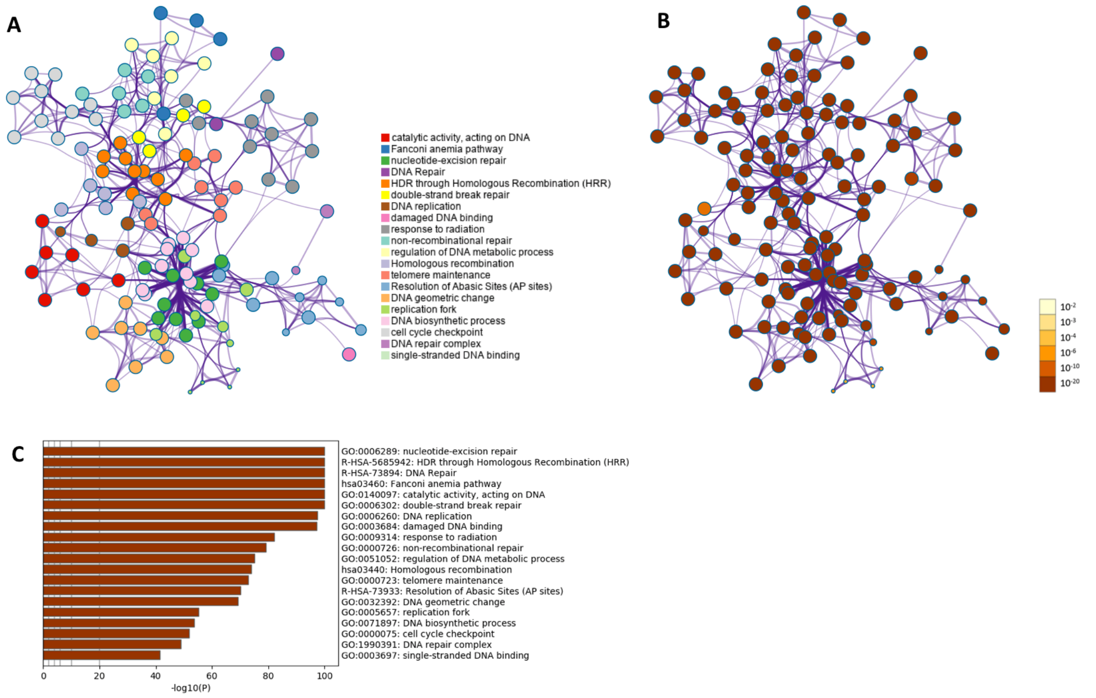


Supplementary Figure 1. **Functions and pathways enrichment analysis provided by Metascape.** The Network represents the top 20 clusters with their representative enriched terms—one per cluster. (A) Each node indicates an enriched term and is coloured by its cluster identity (i.e., nodes of the same colour belong to the same cluster). Node size is proportional to the number of input genes fall into that term. Terms with a similarity score > 0.3 are linked by an edge (the thickness of the edge represents the similarity score). One term from each cluster is selected to have its term description shown as label. (B) The same network but its nodes are colored by p-value, as shown in the legend. The dark the color, the more statistically significant the node is. (C) List of the top 20 statistically enriched terms. The terms can be GO/KEGG terms, canonical pathways, hall mark gene sets, etc.

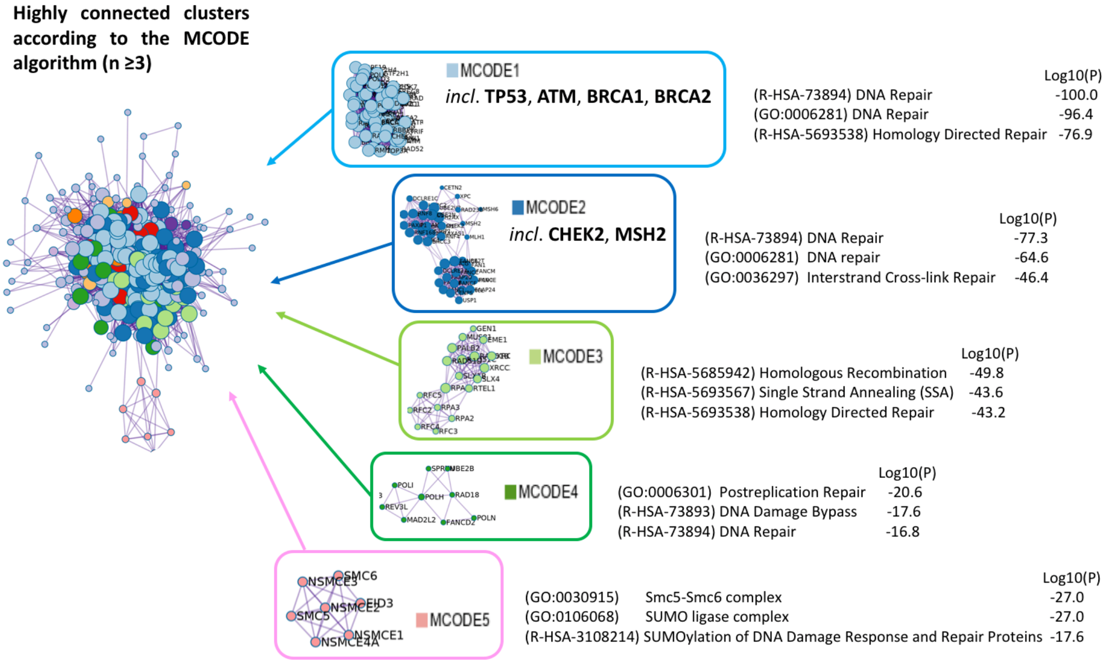


Supplementary Figure 2. **MCODE analysis of the NetDDR network as provided by Metascape**. Densely connected proteins (MCODE clusters) are highlighted. Each cluster is assigned a unique color. GO enrichment analysis was applied to each cluster to assign "meanings" to the network component, where top three best p-value terms were retained.

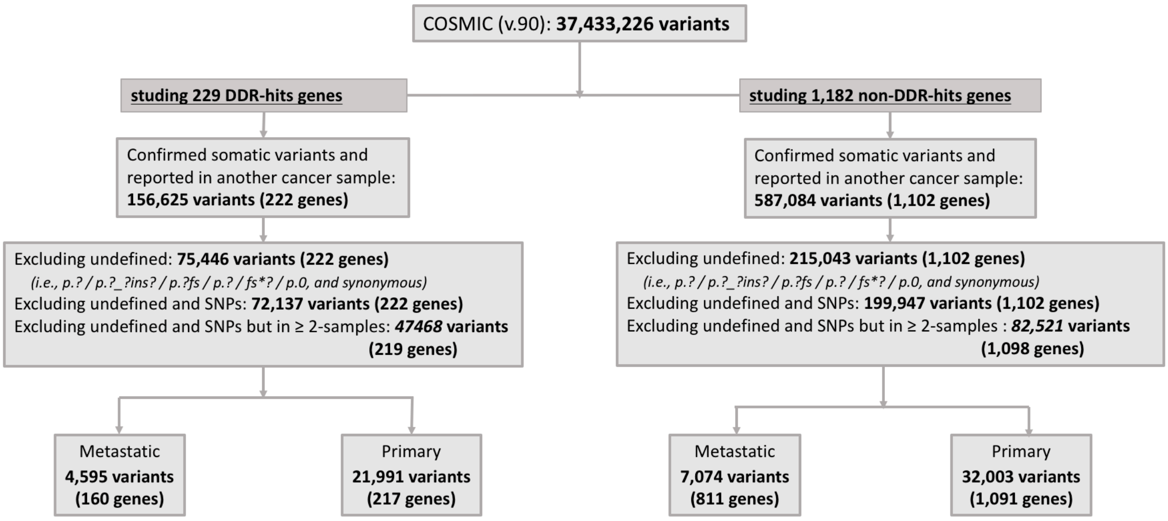


Supplementary Figure 3. **Flow chart for the classification of somatic variants extracted from COSMIC database**. DDR-hits and non-DDR-hits are according to the NetDDR network.

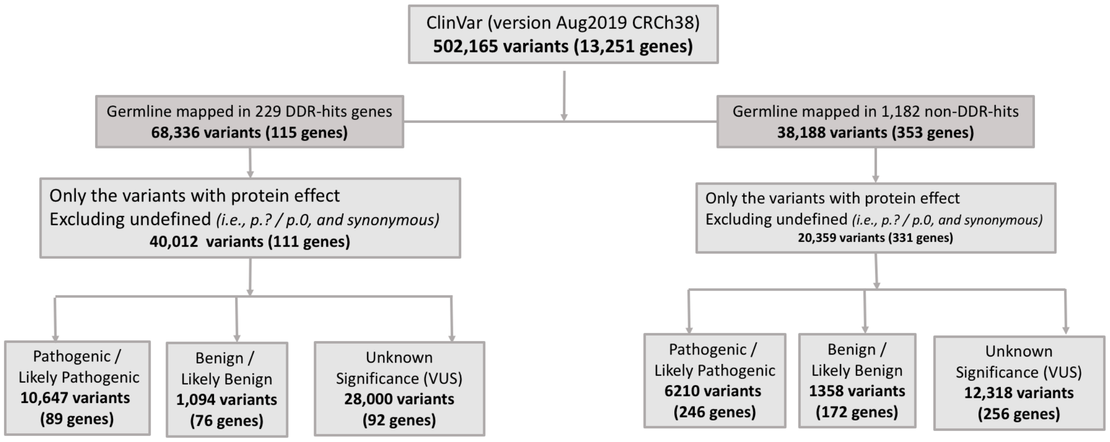


Supplementary Figure 4. **Flow chart for the classification of germline variants extracted from Clinvar database**. DDR-hits and non-DDR-hits are according to the NetDDR network.

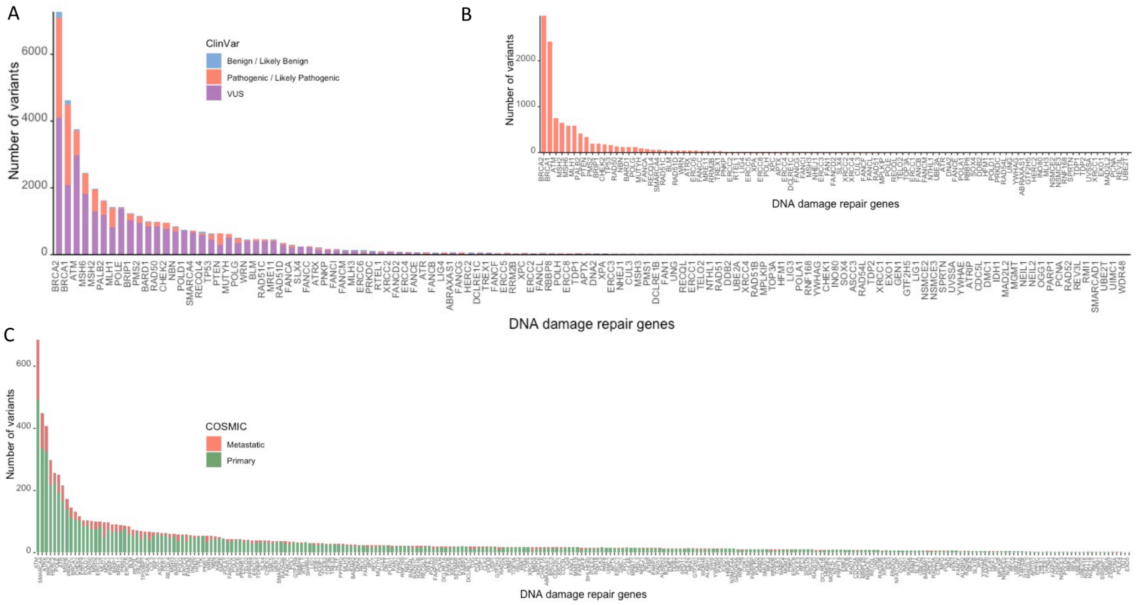


Supplementary Figure 5. **Summary of germline and somatic variants extracted from ClinVar and COSMIC databases.** (A) Barplot of the selected germline variants identified in DDR genes and extracted from the ClinVar database. (B) Subset of pathogenic germline variants (only). (C) Barplot of the selected somatic variants identified in DDR genes and extracted from the COSMIC database. For a better visualization of the barplot, we exclude TP53, IDH1 and PTEN that accumulate a large number of variants.
